# DiffDriver Detects Context-Associated Selection of Cancer Driver Genes

**DOI:** 10.64898/2026.04.06.716771

**Authors:** Jie Zhou, Qirui Zhang, Li Song, Xin He, Siming Zhao

## Abstract

Positive selection of somatic mutations is a driving force for cancer progression. Growing evidence shows that the emergence of a driver mutation in a tumor sample depends on individual-specific contexts, for example environmental exposures or the germline genetic background. Mutations in a driver gene could potentially bring different growth advantages in different contexts, resulting in differential selection of genes in varying contexts. Identifying which contexts modulate selection strength provides critical insights into the selection forces driving tumorigenesis. However, due to the sparsity of somatic mutations and heterogeneous background mutational process across positions and individuals, identification of differential selection has limited power with current statistical tools and is prone to false positives. To address this, we developed a powerful statistical method, DiffDriver, that identifies associations between contexts and selection strength of a driver gene across individuals. DiffDriver accounted for variations of mutation rates across bases and individuals, while taking advantage of functional information of sequences to improve power. In simulations, DiffDriver reduced false positives and boosted power compared to current methods. Multiple individual-level factors created significant heterogeneity in the strength of selection acting on driver genes: 24.3% of driver genes showed differential selection in at least one of the clinical traits studied, and 8.2% of genes had differential selection in different tumor immune subtypes. These results provide insights into the context-associated forces driving cancer evolution.

**Significance:** DiffDriver is a powerful statistical method that identifies associations between individual-level contexts and the strength of selection of driver genes, which revealed a substantial degree of differential selection of cancer genes.

## Introduction

Somatic mutations that confer a growth advantage to cells are under positive selection and are key drivers of cancer^1,2^. Analysis of somatic mutations in tumor patients have identified hundreds of genes under positive selection, or so-called driver genes^3^. Identification of driver genes has been a major focus for large-scale tumor sequencing projects and many computational methods have been developed^4–8^. Such computational methods detect positive selection by comparing the observed mutation rate with the background mutation rate, where an evaluated rate indicates positive selection. Due to the sparsity of somatic mutations, all these methods detect positive selection on aggregated somatic mutations across tumor samples to improve power, assuming selection strength is shared among samples. While such methods have successfully identified many driver genes, individual-specific selection signals are lost due to aggregation of data.

Growing evidence suggests that there are varying environments across individuals and they may put differential selection pressure on some driver genes^9–12^. The driver gene mutation may confer greater selective advantage for the tumor cell in some specific environments than others, leading to stronger positive selection and higher frequency of mutations in patients with these specific environments (**Fig. 1a**). The environment here can be the individual’s genetic background. For example, individuals with a genetic variant in *RBFOX1* altering the splicing pathways show an 8-fold increase of mutation incidence in *SF3B1*, as *SF3B1* mutations work synergistically with the *RBFOX1* variant in promoting cancer progression. Tumor immune microenvironment has also been shown to affect the selection of immune-related driver genes. For example, tumors with immune cell infiltration are enriched with mutations in *B2M* and *HLA* genes, as mutations in these genes help the cells to escape from immune cell cytolytic activities^12,13^. On the other hand, the driver mutation will affect the environment and tumor phenotype, such as tumor microenvironment or tumor progression process. For example, selection on certain driver mutations has been found specific to the metastatic stage of tumor^14^. Environmental exposures likewise shape selection: in lung cancer, tobacco smoking and airborne particulate pollution are associated with distinct driver landscapes—for example, airborne fine-particulate pollution promotes selection of EGFR-mutant lung adenocarcinoma^10^. These examples motivate us to more systematically identify associations between a variable context and the selection strength of driver genes. We refer to driver genes showing such association with a potential context as genes under “differential selection” or “differentially selected” genes. In contrast, genes whose selection strength do not associate with the context variable are referred to as the constitutively selected genes.

**Fig. 1.**
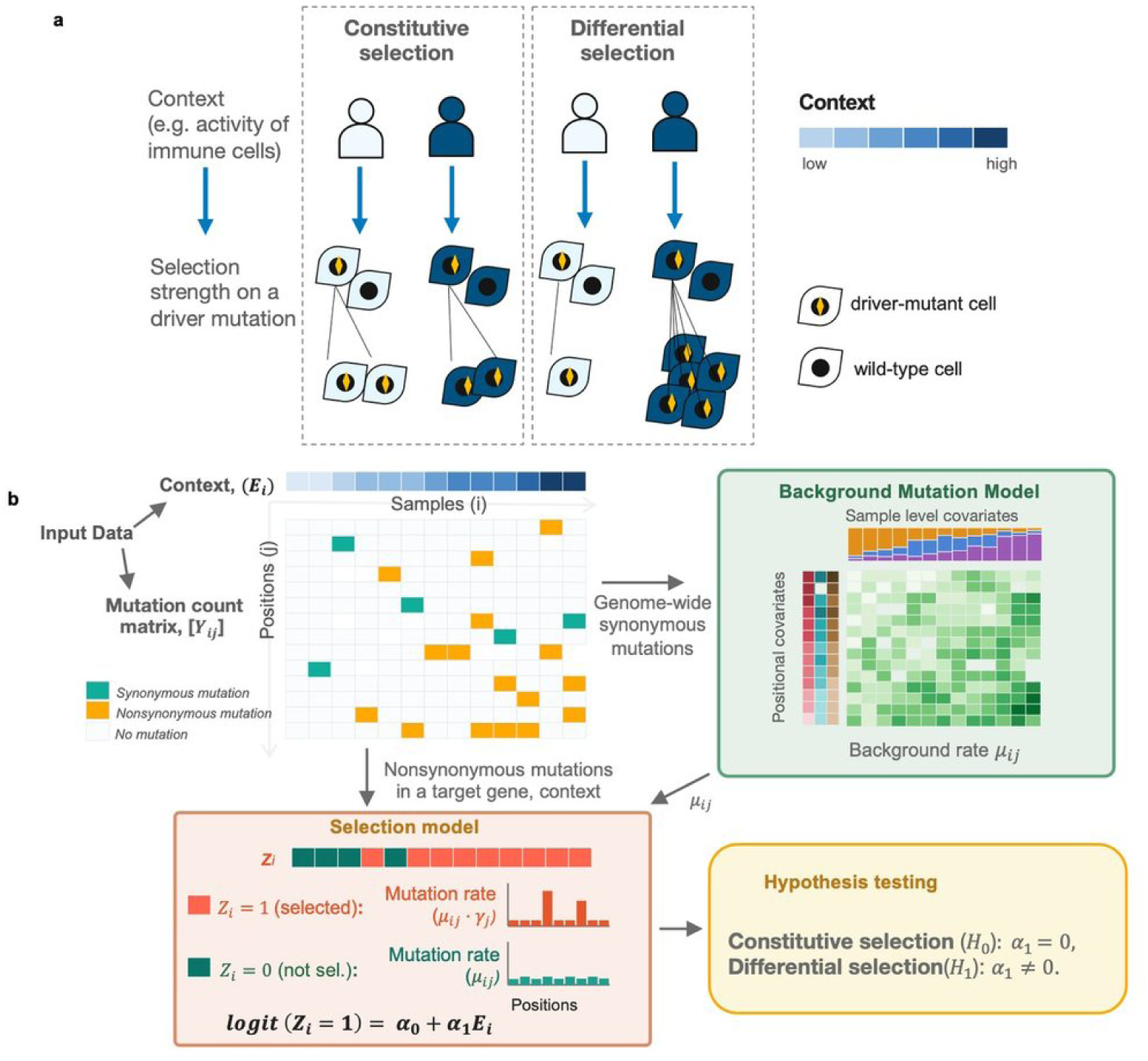
The DiffDriver framework for detecting context-associated selection on cancer driver genes. **a**, Concept. DiffDriver tests whether an individual-level "context" (*E_i_* for sample *i*; e.g., activity of immune cells) modulates the strength of positive selection on a driver gene. Under constitutive selection, a driver mutation confers the same growth advantage regardless of context, so the frequency of driver-mutant cells is similar across individuals; under differential selection, the advantage—and hence the frequency of driver-mutant cells—varies with the context. The blue color scale denotes the context value from low to high; driver mutations conferring growth advantage is labeled as a yellow diamond. **b**, Model schematic. Input data of DiffDriver are a matrix indicating the presence of somatic mutations over all individuals and positions ([*Y_ij_*]), as well as matching per-individual context, (*E_i_*). Genome-wide synonymous mutations are used to train the background mutation model (BMM, green box), which estimates the position- and individual-specific background mutation rate *μ_ij_* (green matrix). It learns individual specific rates by decomposing the mutational process into a small number of mutational signatures and obtain per-sample signature loadings (top), it then accounts positional covariates, such as replication timing and expression and a gene specific effect learned from data(left). Nonsynonymous mutations in a target gene are then modelled by the selection model (orange box). A latent indicator *Z_i_* denotes whether the gene is under selection in individual *i*. When the gene is not under selection (*Z_i_* = 0) mutations occur at the background rate *μ_ij_*; when under selection (*Z_i_* = 1) the rate is elevated to *μ_ij_γ_ij_*, where *γ_ij_* scales with functional annotations of the mutation. DiffDriver then tests the context coefficient (*α*_1_) to distinguish differential selection (*H*_1_: *α*_1_ ≠ 0 ) from the null hypothesis of constitutive selection (*H*_0_: *α*_1_ = 0 ) (yellow box).

In this study, we developed a general statistical framework to identify genes under differential selection with respect to a context. Conceptually, this is similar to genome-wide association studies (GWAS)^15^ that explore genotype-phenotype relationship, and is often done via regression analysis. Applying such methods on the differential selection problem means to simply correlate the number of somatic mutations in each gene with the context variable. Indeed, these methods and other standard association tests are what current studies use to solve this problem^12,16–18^. For example, Fisher’s exact tests have been used to identify driver genes affected by germline background^16^, linear-regression–based methods to identify drivers enriched in different immune contexts^13,18^, and and the Mann–Whitney test to call metastasis-enriched driver genes relative to primary tumors^14^. However, such standard association testing methods have severe limitations when they are applied to detecting differential selection. The challenges come from two aspects: 1, due to the sparsity of somatic mutations, these methods often have limited power; 2, background mutation rates vary across patients and positions, confounding the relationship of selection strength and context variables and resulting in potential false positives. To address these challenges, we propose a model-based approach, called DiffDriver, to identify genes under differential selection. DiffDriver models background mutation rate and strength of positive selection at per-sample, per-base-pair level. We demonstrate in simulations that DiffDriver effectively controls for false positive rate as it carefully models background mutation rate. It also improves power by leveraging various functional annotations of mutations to capture selection signals.

We applied DiffDriver to a number of potential contexts that may lead to differential selection, including the patient’s age, tumor’s genomic background, immune environment, etc. Our results show that selection is often heterogeneous, with 24.3% of cancer drivers gene exhibiting differential selection in at least one clinical context tested in this study. We found a number of driver genes, such as *KRAS* and *TP53*, exhibiting differential selection under different immune sub-types, highlighting the potential of these genes in modulating tumor-immune interactions. We believe the application of DiffDriver on large-cohort of cancer patients somatic mutation data will be critical for understanding the individual-specific selection of driver genes and cancer progression.

## Materials and Methods

### The DiffDriver background mutation model

A key challenge in detecting differential selection is accurately estimating background mutation rates at the individual and position level. Somatic mutations arise from diverse mutational processes—including replication errors, DNA damage from environmental exposures, and deficiencies in DNA repair—that vary substantially across individuals and genomic positions. If these differences are not properly modeled, they can confound the relationship between selection strength and the context variable, producing spurious associations. Prior driver-detection methods—notably MutSigCV^4^ and dNdScv^8^—also model patient-and covariate-level heterogeneity in the background mutation rate; DiffDriver builds on this idea and further resolves the rate to the per-sample, per-position level required for differential-selection testing. The DiffDriver background mutation model (BMM) addresses this challenge in two steps. First, we estimate the expected mutation rate for each type of mutations in each sample, where a mutation type is defined by both the substitution and the flanking bases. This step accounts for sample-specific mutational features by decomposing the observed mutation spectrum into latent mutational signatures using topic modeling, which captures the sample-specific composition of active mutational processes (e.g., APOBEC-mediated mutagenesis, defective mismatch repair). Each sample receives a personalized mixture of these signatures, yielding a sample- and mutation-type-specific baseline rate. Second, we adjust this baseline rate for position-specific genomic covariates that are known to influence local mutation rates, such as replication timing, gene expression level, and CpG context^4^. Together, these two steps produce a position- and individual-specific background mutation rate that reflects the true baseline expectation in the absence of selection.

In more detail, our BMM first models the mutation rate of any type *t* in a sample *i*, denoted as *λ_it_*, and then adjusts for position-specific covariates. To model *λ_it_*, we use a topic modeling approach similarly as described before^19^. The key idea is that *λ_it_* is determined by the total number of mutations in sample *i* (*N_i_*) and the sample-specific composition of mutational signatures. Topic modeling factorizes the silent mutation count matrix across samples into sample-specific loadings *l_ik_* (the contribution of signature *k* in sample *i*) and signature-specific factors *f_tk_* (the relative rate of mutation type *t* in signature *k*), borrowing strength across all samples to reliably estimate per-sample rates. The loadings and factors are jointly estimated from silent mutation counts using the fastTopics package, which performs non-negative matrix factorization. We have:

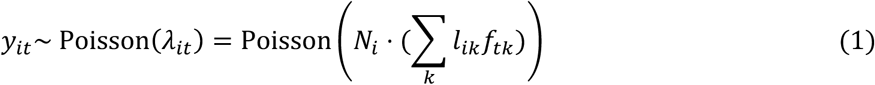

where *y_it_* is the total number of mutations of type *t* in sample *i*.

Once *λ_it_* is estimated, we adjust for position-specific covariates *X*_j_ (e.g., replication timing, gene expression, CpG context). Using a Poisson log-linear approach, the deviation of the mutation rate at position *j* of type *t* from the expected rate *λ_it_* is given by *e^X^*^j^*^β^*, where *β* is the effect-size vector. Because the sum of mutation rates of type *t* across all positions in sample *i* must equal *π_it_*, we normalize by *a_t_* = ∑_j_ *e^X^*^j^*^β^*. Putting these together, the per-position mutation count *y_ij_* is modeled as *y_ij_* ∼ Poisson(*μ_ij_*), with:

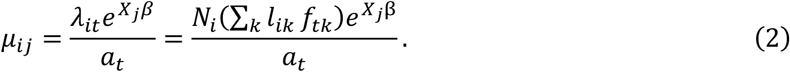

The effect sizes *β* are shared across samples and estimated from silent mutations by maximum likelihood, as described in the driverMAPS paper^5^.

As a comparison, we also constructed a version of the BMM without adjusting for mutational signatures, called basic BMM. In this version, we have 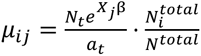, where *N_t_* is the total number of silent mutations with type *t*, 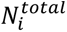 is the number of silent mutations in sample *i*, *N^total^* is the total number of silent mutations in the cohort.

### The DiffDriver selection model

The selection model builds on the background mutation rates estimated by the BMM and aims to detect whether the strength of positive selection on a driver gene varies with an individual-level context variable. The core idea is to introduce a latent binary indicator for each individual that represents whether the gene is under selection in that individual’s tumor. Whether selection is “on” or “off” is modeled as a function of the context through logistic regression, linking individual-level characteristics to selection probability. When selection is active, the observed mutation rate deviates from the background rate, and the magnitude of this deviation at each position is informed by functional annotations of the mutation—such as conservation, predicted deleteriousness, and hotspot status.

We model selection given background mutation rate to depend on an individual level variable (*E_i_*) and use it to fit the nonsynonymous mutation data of a gene. The first component of the model determines whether a gene is under selection in a given individual. We introduce a latent variable *Z_i_* ∈ {0,1} to indicate if the gene is under selection in sample *i*, we have

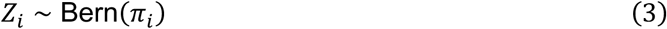

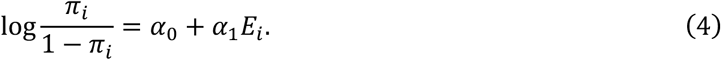

At a single base position, a mutation either occurs or not in a given individual, so the per-position, per-individual indicator is naturally modeled with a Bernoulli distribution in the selection step. This is consistent with the Poisson used for genome-wide background-rate estimation, which aggregates counts over many positions and mutation types, because at the very low per-position rate a Poisson count reduces to a 0/1 Bernoulli outcome.

Equations (3) and (4) together model the probability that a gene is under selection in a given individual as a function of the context. Equation (3) introduces the latent selection indicator, and Equation (4) links it to the context variable through a logistic regression, where the coefficient α₁ captures the effect of the context on the probability of selection. The second component of the model specifies how mutation counts are generated conditional on whether the gene is under selection:

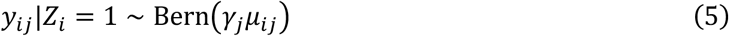

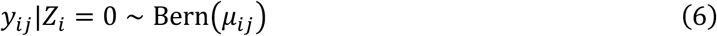

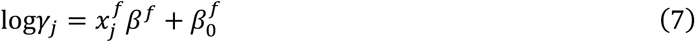

Equations (5) and (6) specify the mutation likelihood under the two selection states. When the gene is under selection in individual *i* (Equation 5), the mutation rate at mutation *j* is modified by *γ*_j_ , which represents the selection effect. When the gene is not under selection (Equation 6), mutations arise at the background rate alone. Equation (7) models the selection effect *γ*_j_ as a log-linear function of functional annotations at mutation *j*, such that mutations with stronger predicted functional impact (e.g., nonsense mutations, conserved sites) receive greater deviations from the background rate.

Under the null model, *π_i_* does not depend on *E_i_*, we have *α*_1_ = 0. Under the alternative model, which means the gene is under differential selection, *α*_1_ ≠ 0.

### Adjusting for confounding covariates

The association between a context and selection strength can be confounded by other individual-level variables. To account for this, DiffDriver can incorporate additional covariates into the logistic regression that determines the selection probability (Equation 4). Denoting the covariate vector for sample *i* as *w_i_*, the log-odds of selection become *α*_0_ + *α*_1_*x_i_* + *γ^T^w_i_*, where *x_i_* is the context of interest and *γ* are the covariate effect sizes. The covariates are included in both the null (*α*_1_ = 0) and the alternative (*α*_1_ ≠ 0) models. We recommend adjusting for only a small number of covariates: including many can prevent the logistic regression from converging or yield unstable estimates within the EM algorithm, particularly for genes with few mutated samples. In all real-data analyses in this study, we adjusted for patient age and sex, except for the female/male-specific cohort (e.g. UCEC), for which we adjusted for age only.

### Parameter inference and model selection

In the selection model described above, the site-specific mutation rate, *μ_ij_*, is estimated using the Background Mutation Model (BMM) and treated as a fixed parameter. The parameter 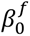 represents the average selection strength across all positions within a gene and is estimated directly by our model. The vector *β^f^* denotes the effect sizes associated with specific functional annotations.

For the functional effect sizes, we utilized values of *β^f^* previously estimated by DriverMAPS^5^, which aggregates mutation counts across cohorts under the assumption that all samples are subject to selection for a given driver gene. Because DriverMAPS averages effects across all samples, the resulting *β* estimates are likely lower than the true effect sizes expected in the context-specific DiffDriver model. However, we intentionally adopted these DriverMAPS-derived parameters to ensure a conservative estimation of differential selection, thereby reducing the likelihood of false positives. In a sensitivity analysis, using the imported, averaged functional effect sizes rather than re-estimating the true, context-specific effect sizes reduced DiffDriver’s power by less than 2% across all sample sizes, while the false discovery rate remained controlled (Supplementary Figure 10).

We use an EM algorithm^20^ to estimate parameters *θ*= {*α*_1_, *α*_0_, 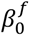} (see Supplementary Information).

### Simulations

We randomly select a gene (*ERBB3*) to simulate mutations and test for differential selection of this gene. For the null simulation (**Fig. 2**), we simulated genome-wide synonymous mutations under the DiffDriver BMM. We set *f_tk_* and *l_ik_* to those learned from TCGA LUAD data (total mutational signatures = 6, mutation types = 96). We simulate a binary context (*E_i_*) to correlate with the loadings of one of the mutational signature (*l*_1_). We simulated nonsilent mutation data for this gene under the DiffDriver selection model for an oncogene with two hotspots. We set *α*_1_ = 0, i.e. no differential selection for the null simulation. The other parameters (coefficients for functional and mutational annotations) were set to those learned from the TCGA LUAD data as in driverMAPS. The selection strength (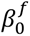) is set to the average strength of 53 oncogenes^2^ across all samples. We ran DiffDriver and Coselens on the simulated genome-wide synonymous mutations and nonsynonymous mutations in this particular gene. For multiple linear regression, we adjust the linear model using total number of genome-wide synonymous mutations. For other methods, nonsynonymous mutations in this particular gene is used. We performed this simulation 100 times to obtain the false positive rate for each method.

**Fig. 2.**
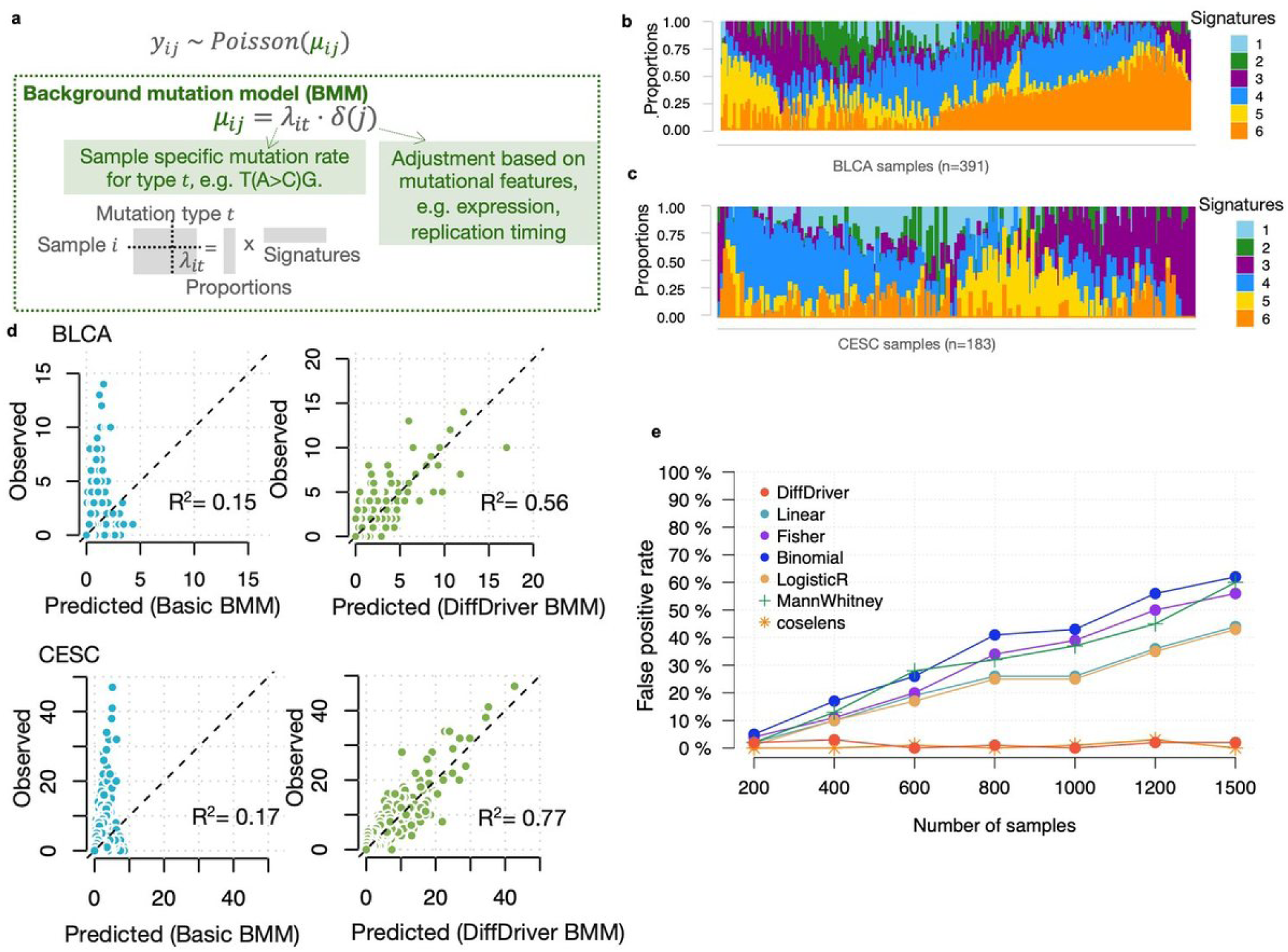
The DiffDriver background mutation model (BMM) captures individual- and position-specific background mutation rates. **a,** Schematic of the BMM. The somatic mutation count at position *j* in sample *i*, *y_ij_*, is modeled as *y_ij_* ∼ Poisson(*μ_ij_*), where the background rate *μ_ij_* = *λ_it_* ⋅ *δ*(*j*). *λ_it_* is the background mutation rate for mutation type *t*, (defined by the substitution and its trinucleotide context) in sample *i*, estimated by topic modeling, which decomposes each sample’s mutation spectrum into a mixture of mutational signatures—the per-sample signature proportions (loadings) times the signature profiles. *δ*(*j*) further adjusts the rate for position-level features such as gene expression and replication timing and a gene specific effect derived from data (see Methods.) **b, c,** Per-sample composition of the six mutational signatures inferred by the BMM, for (b) bladder cancer and (c) cervical cancer (CESC). Each vertical bar is one sample; segments give the proportion (loading) of each signature and sum to one. Signatures 1–6 are learned de novo by topic modeling of the 96 trinucleotide mutation types (shown in Supplementary Fig. 2). Samples are ordered based on similarity of their loadings as described in a structure plot^48^. **d,** Goodness-of-fit of the BMM. For BLCA (top) and CESC (bottom), observed versus predicted mutation counts (per trinucleotide type, on held-out chromosomes) are shown for the Basic BMM (left; mutation type and overall sample rate only) and the DiffDriver BMM (right; additionally modeling inter-individual signature composition). The dashed line denotes y = x, and the coefficient of determination (R²) is indicated. **e,** DiffDriver controls the false positive rate in a null simulation. In this simulation with no differential selection, the loading of one mutational signature (e.g., mutations resembling oxoG artifacts) is correlated with the context. DiffDriver estimates the background rate directly from simulated genome-wide silent mutations (see Methods for details).

To test the power of DiffDriver and comparators (**Fig. 3**), we simulate silent mutations in 200 randomly selected genes using mutation type specific rate averaged cross tumor types, without mutational signatures. We simulate nonsilent mutations for these genes under the DiffDriver selection model as tumor suppressor genes or oncogenes, by setting different *β^f^*. For all our simulations, we simulate half of samples under selection so we set *β^f^* as 2 fold of the values as learned in drvierMAPS, which is averaged across all samples.

**Fig. 3.**
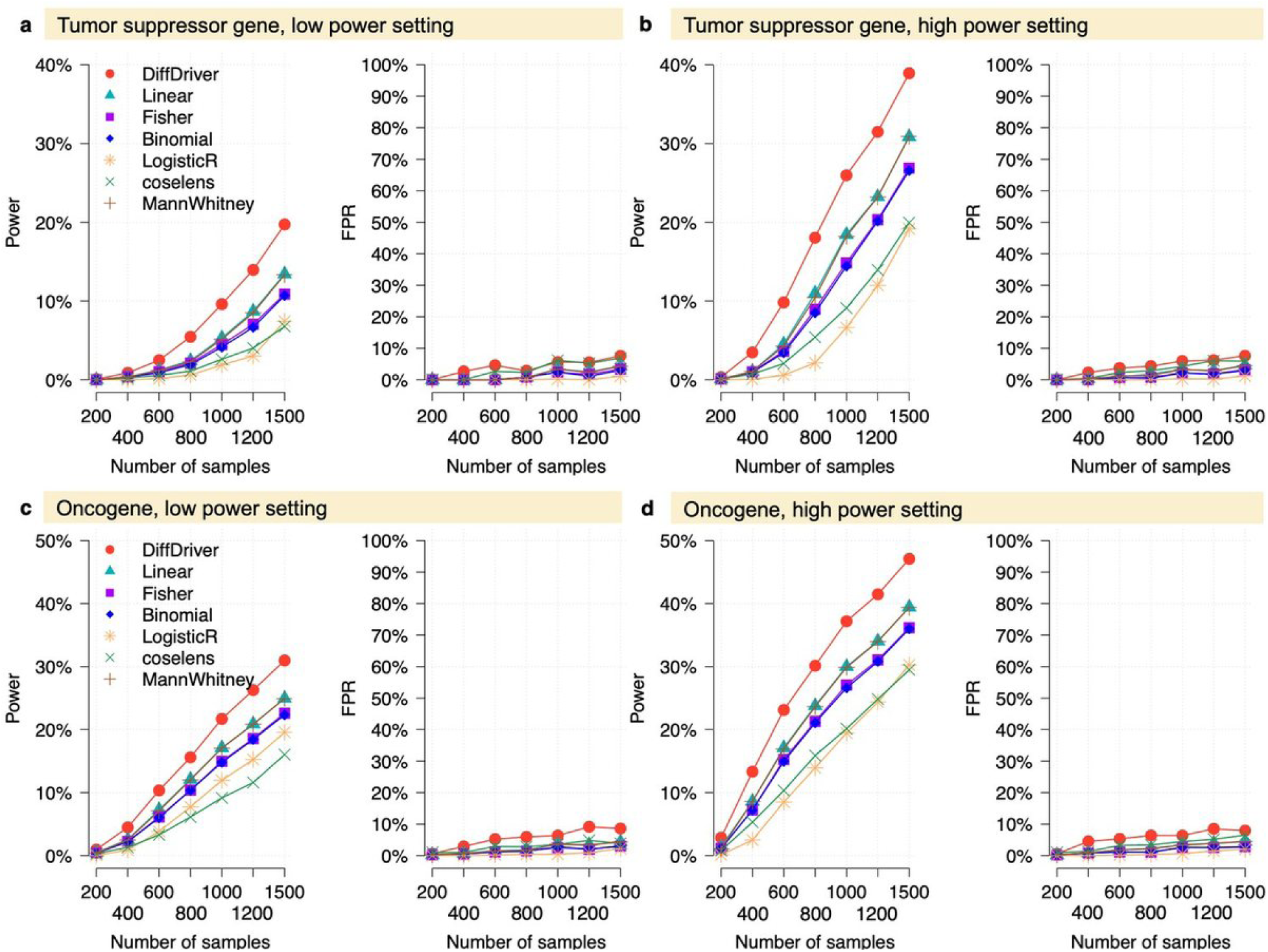
Benchmarking DiffDriver performance against standard methods using simulated data. **a–d,** Performance comparison of DiffDriver (red) with four baseline methods (Linear regression, Fisher’s exact test, Binomial test, Logistic regression, coselens^17^ and Mann Whitney test) in detecting driver genes. The simulations evaluate performance for (**a, b**) tumor suppressor genes and (**c, d**) oncogenes under low-power (**a, c**) and high-power (**b, d**) settings. The x-axis represents the cohort size (number of samples). In each panel, the left plot shows the statistical power, and the right plot indicates the False Positive Rate (FPR).

We set different *α*_1_ and *α*_0_ to simulate the individual level context (*E_i_*). Of the 200 genes, we simulate context (*E_i_*) for 150 genes with *α*_1_=0, which means they were not under differential selection (null genes); for the rest 50 genes, we simulate with nonzero *α*_1_ which means they are under differential selection. We evaluated power as the fraction of the genes that pass a significance cut off (Benjamini–Hochberg FDR = 0.1) in the 50 true differentially selected genes and false positive rate as the fraction of null genes that pass the significance cut off. We benchmarked our model performance and a few comparator methods in several simulation settings: 1. Simulation of a binary context. In the high power setting, we set *α*_1_ and *α*_0_ so that the fraction of samples under selection is 10% and 90% in the two groups with different context values, in the low power setting, we set the fraction of samples under selection to 20% and 80% in the two groups. 2. Simulation of a continuous context. We simulate *E_i_* using a normal distribution (*N*(0, 2^2^). In the high-power setting, we set *α*_1_ and *α*_0_ so that the sample with 75th percentile context value is 10x more likely under selection then a sample with 25th context value. In the low power setting, this ratio is 3.5x. For each setting, we evaluated the power and false discovery rate for tumor suppressor genes or oncogenes.

### Application of DiffDriver on TCGA data

We used somatic mutations from the 20 tumor types used in Zhao et al, 2019^5^ . The tumor types and their abbreviations: bladder cancer (BLCA), breast cancer (BRCA), cervical squamous cell carcinoma and endocervical adenocarcinoma (CESC), cholangiocarcinoma (CHOL), esophageal carcinoma (ESCA), glioblastoma multiforme (GBM), head and neck squamous cell carcinoma (HNSC), kidney chromophobe (KICH), kidney renal clear cell carcinoma (KIRC), kidney renal papillary cell carcinoma (KIRP), liver hepatocellular carcinoma (LIHC), lung adenocarcinoma (LUAD), lung squamous cell carcinoma (LUSC), pancreatic adenocarcinoma (PAAD), prostate adenocarcinoma (PRAD), sarcoma (SARC), skin cutaneous melanoma (SKCM), testicular germ cell tumors (TGCT), uterine corpus endometrial carcinoma (UCEC), uterine carcinosarcoma (UCS) .

We obtained clinical traits of the samples (FGA, TMB, patient age, disease-specific survival, progression-free survival) from the TCGA data portal (https://portal.gdc.cancer.gov/). We calculated immune repertoire diversity of TCGA samples using the data generated by TRUST4^21,22^. Specifically, we defined entropy for a receptor chain in a sample as 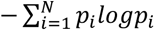 *p_i_logp_i_*, and clonality as 1 − *entropy*/*log N*, where N is the number of distinct clonotypes, i.e., CDR3 nucleotide sequence, and *p_i_* is the abundance fraction of clonotype *i*. We run DiffDriver for each tumor type for genes showing positive selection, as identified in Zhao et al, 2019. When running DiffDriver, we adjusted for age and sex (except for unisex tumor types). For pan-cancer summaries, we pooled tests across tumor types and applied the Benjamini–Hochberg (BH) procedure. For results in individual tumor types, we applied BH to tests within the tumor type. Immune subtype is treated as a context variable in exactly the same way as any other context. Because it is categorical (C1-C4), we encode it as one-vs-rest binary indicators (e.g., C2 vs. all other subtypes). There are originally 6 subtypes from ref^12^, C5 and C6 has < 5% samples in most cohorts analyzed in this study, thus we excluded them. We ran DiffDriver on each context separately.

### Pan-cancer immune-subtype analysis

To characterize differential selection across immune subtypes at the pan-cancer level, we pooled tumor samples across cancer types rather than analyzing each tumor type separately. Starting from the 299 driver genes of Bailey et al.^3^, we used point mutations (single-base substitutions) from the TCGA PanCancer Atlas and excluded hypermutated samples (>1,000 mutations per sample). For each gene we counted point mutations per tumor type, retained the tumor types with at least 5 point mutations, and kept genes with more than 20 point mutations in total. Each gene’s cohort was defined as the union of its qualifying tumor types among the 27 cancer types carrying immune-subtype labels, and genes sharing the same set of tumor types were grouped so that the background mutation model was fit once per cohort. This ended with 220 genes that we ran DiffDriver on. Each gene was tested for differential selection between each immune subtype and all remaining subtypes (Thorsson C1–C4, one-vs-rest), adjusting for patient age, sex, and tumor type; sex was dropped for single-sex cohorts and tumor type for single-tumor cohorts. Because the pooled cohorts span a wide range of sizes, we scaled the number of mutational signatures *k* used in the background mutation model to cohort size: *k* = 6 for cohorts of 1,000 or fewer samples, increased by 1 for every additional 200 samples above 1,000, and capped at *k* = 20. Significance was assessed from the DiffDriver likelihood-ratio test p value, and the Benjamini–Hochberg false discovery rate was computed separately across all genes within each immune subtype. Gene–immune subtype associations with FDR < 0.1 were considered significant.

## Results

### A probabilistic model to test for differential selection with a context

Our method, DiffDriver, tests for an association between an individual-level variable (which is a quantitative measure of a “context” of interest) and selection strength on somatic mutations in a driver gene (**Fig. 1b**). The context is an individual-level variable that can potentially drive differential selection, such as the presence or absence of a germline variant, or a phenotype of the tumor that is affected by selection, such as tumor growth rate. We note that for many individual-level measurements of the tumor, it is not easy to distinguish whether they serve as the cause or are a result of differential selection. For example, the tumor immune microenvironment drives somatic mutation selection and can be altered by driver mutations. DiffDriver only tests the association between the given context and selection and does not infer the direction of effect. It also does not infer causality, which is a limitation shared by all association tests. The differentially selected genes are called when their selection strength associates with the context variable.

We give an overview of the DiffDriver model. The model can be viewed as a generalization of the commonly used dN/dS analysis for studying selection in driver genes, which compares the mutation rates in non-synomous vs. synomous mutations. This analysis, however, treats all non-synonymous mutations equally. Our idea is that, in driver genes, mutations are more likely to occur at the likely functional sites. This motivates a model of the mutation count at each position. This count depends on: (1) the "background mutation rate," estimated by a Background Mutation Model (BMM) using genome-wide synonymous mutation data. The background mutation rate has been shown to be substantially heterogeneous across positions and samples^4,8^, and the BMM in DiffDriver accounts for positional and sample-level covariates to obtain a position-and sample-specific rate; and (2) the strength of selection at that position, defined as the deviation of the mutation rate from the background rate, which depends on the functional effect of the mutation. Lastly, to capture context-associated selection, we introduce an indicator variable of whether the gene is under selection in a particular individual. Whether this indicator equals 1 (selection) depends on the specific context of that individual.

Specifically, DiffDriver models mutation count data (*Y*) for each position in a gene across a cohort of individuals. Given the mutation rate at individual *i*, position *j*, the mutation count *Y_ij_* ∈ {0, 1} follows a Bernoulli distribution. We introduce an individual-level latent variable *Z_i_* to indicate if the gene in a particular individual *i* is under selection. When the individual is not under selection (*Z_i_* = 0), the mutation rate in this individual is the same as the background rate (*μ_ij_*). When the individual is under selection (*Z_i_* = 1), the mutation rate in this individual deviate from the background rate (*μ_ij_γ_ij_*), where the parameter *γ_ij_* indicates selection strength and it depends on base-pair level functional annotations. These functional annotations, e.g. mutation consequences (nonsense/missense) and evolutionary conservation, serve as proxies for the likely deleterious or functional impact of each somatic variant.

We assume the latent variable *Z* depends on the context *E* through a logistic regression. When the gene is under constitutive selection with respect to the context (hypothesis *H*_0_), the probability of selection across different individuals is the same (the regression coefficient *α*_1_ = 0); when the gene is under differential selection (hypothesis *H*_1_), the context will affect the probability of selection in an individual (*α*_1_ ≠ 0). DiffDriver will perform hypothesis testing for a driver gene given the mutation count data for this gene, the background mutation rate (*μ_ij_*) and selection rate (*μ_ij_γ_ij_*).

We provided some intuitions of the advantages of DiffDriver in detecting context-associated selection. A number of individual factors have been found to affect mutational signatures composition and thus affect driver mutation distribution across individuals, confounding the standard association analysis and results in false positives. DiffDriver solves this issue by explicitly model mutation rate and selection at per-position per-individual level, accounting for various factors affecting the mutational and selection process. DiffDriver also incorporates functional annotation in modeling the strength of selection, improving the power. Intuitively, say for a binary context, standard association analysis only compares the total number of mutations of the gene in the two context groups, thus given the low mutation numbers, the power is often limited. However, if the mutations in one group are more functional relevant, e.g. nonsense mutations or mutations at conserved sequences, while mutations in the other group are predicted to have low functional impact, we know the selection strength in these groups is likely different, even though the difference in total number of mutations may not be obvious.

### DiffDriver models position and individual specific mutation rate to reduce false positives

To fully account for the various positional and individual factors that affect the mutation rate, the BMM in DiffDriver estimates a position- and individual-specific mutation rate, which is then plugged into the selection model. BMM models the background mutation rate *μ_ij_* in individual *i* at position j with two components (**Fig. 2a**). The first component is parameter *λ_it_*, which is the background mutation rate for nucleotide change type *t* in individual *i*. When implementing DiffDriver, we consider 96 nucleotide change types based on the mutation and its tri-nucleotide contexts (see Methods). Because somatic mutation is very sparse, it is often unreliable to use the data from individual *i* alone to estimate *λ_it_*. To solve this problem, we use topic modeling to model background mutation count data across all samples and all nucleotide change types (see Methods). In this way, DiffDriver captures per-sample bias—including both biological mutational signatures, such as smoking-related signatures, and technical artifacts, such as platform-specific spectra—directly from the data, without requiring these factors to be known or measured in advance. We further modify *λ_it_* based on the gene’s features that could affect mutation rate, such as expression, replication timing etc. It also incorporated a random effect to learn gene specific mutation rate (see Methods). We use synonymous mutation data to learn parameters in BMM, under the assumption that the vast majority of synonymous mutations are not under selection.

We found that the BMM of DiffDriver can capture the heterogeneity of background mutation rate across individuals. We show two example results from applying DiffDriver on The Cancer Genome Atlas (TCGA) data (Fig. 2b-d show results for bladder cancer, BLCA and cervical cancer, CESC). CESC contains 183 samples and an average of 25.57silent mutations per sample, BLCA contains 391 samples and average of an average of 62.21 silent mutations per sample. We found that for most cancer types, the mean square error of the BMM drops the fastest with an increasing number of “mutational signatures” before 6 (Supplementary Figure 1). We thus used 6 mutational signatures as the default (see Methods). DiffDriver estimates the proportion of “mutational signatures” for each individual and we see it varies widely from sample to sample (**Fig. 2b-c**). The “mutational signatures” identified by DiffDriver capture APOBEC3 related mutational processes which has been reported as the most dominant mutational processes for these two types of cancers (featured by C>T and C>G mutations in TCW motifs) (Supplementary Figure 2).

We then compared the performance of the BMM of DiffDriver and a mutation rate model (Basic BMM) that accounts for mutation type and overall sample rate difference but not interindividual mutational signature difference (**Fig. 2d**). The Basic BMM aggregates mutations across samples to get mutation rate for each nucleotide-change type and then adjusts it based on the total number of mutations in the sample to obtain sample specific rate (see Methods). We trained DiffDriver BMM and Basic BMM using mutation data from the even number chromosomes (chr2, 4, …, 22) and made predictions on the remaining chromosomes for each nucleotide-change type (96 total) in each sample. The predicted values from DiffDriver showed better correlation for all tumor types than Basic BMM.

We then benchmarked DiffDriver against several commonly used association methods on simulated data with heterogeneous background mutation rate. We simulated a binary context which did not affect selection of the driver gene but affected the proportions of mutational signatures (See Methods) across samples. Under this scenario, no differential selection should be detected. For linear and logistic regression, we regressed the context against mutation count in the driver gene, using total mutation count from the sample as a covariate. For the Mann-Whitney, Fisher and Binomial tests, we binarized the context and tested the difference of mutation count in the driver gene in two groups of samples with different contexts. We also tested Coselens^17^, which compares the strength of selection (using the dNdScv framework) on a gene between two predefined groups of samples. We therefore also binarized the context for Coselens. Coselens estimates a group-level background mutation rate and this will help to adjust the confounding mutational signature. We see DiffDriver and Coselens’ false positive rate remains low with increasing cohort sizes, but the other methods are inflated with false positives (**Fig. 2e**). We further verified that DiffDriver’s calibration is robust across data settings by varying the generative assumptions (Supplementary Fig. 4). As the context–signature confounding increases (R² = 0.3→0.95), count-based methods’ false-positive rate rises monotonically (≈10%→≈60%), whereas DiffDriver and Coselens remain controlled (Supplementary Fig. 4a).

This control is insensitive to mutation-burden heterogeneity (a ≈5–10× per-sample spread) (Supplementary Fig. 4b); to context type, holding for a continuous confounded phenotype (Supplementary Fig. 4c); and to the number of fitted signatures (k ∈ {4–10}, Supplementary Fig. 4d). Thus, DiffDriver ensures calibrated results to identify differential selection under a context, even when the context affects the background mutational process in a sample-specific way.

### DiffDriver leverages functional annotations to model selection and improves power

DiffDriver models selection as the deviation of observed mutation rate from the background mutation rate and is denoted as *γ_ij_* for position *i*, sample *j* (**Fig. 1b**). We reasoned that selection strength on the position depends on its functional importance and incorporating functional annotation of mutations can improve power to detect selection. For example, consider the case where we are testing differential selection of a tumor suppressor gene (TSG) with a binary context. If we see many more nonsense mutations in one context group than the other group, then it means selection is likely stronger in the former group, even though the total mutations in this gene may be similar in these two groups. Therefore, we model *γ_ij_* to depend on the functional annotations of the mutation, including conservation, whether it is a nonsense or splice-site mutation, PhyloP, SiFT and GERP scores (see Methods). As another annotation, we consider whether a mutation is located in a “hotspot”, which represents a particularly important functional site/region of a driver gene. To define these hotspots, we used the aggregated mutation count data across individuals as previously described^5^. We included the hotspot status of mutations as an annotation for well-known oncogenes previously curated^2^. The effect sizes for each annotation are learned from real data using DriverMAPS^5^ and separately for TSGs and oncogenes (see Methods).

We benchmarked the power and false positive rate of DiffDriver and other methods through extensive simulations. In this simulation, we assumed a uniform background rate for all mutation types across individuals to eliminate background mutational effects. Consequently, the results reflect each method’s performance based purely on its ability to detect differential selection, independent of how the background mutation rate is estimated. We simulated selection to depend on functional annotation using realistic parameters estimated from real data (see Methods). We first tested the scenarios when the context is binary, essentially there are two groups for this cohort, and the probability of an individual under selection is different in this cohort. We simulate the selection status (*Z_i_*) of individual *i* to follow a logistic regression depending on the context (*logit*(*P*(*Z_i_* = 1)) = *α*_1_ *E_i_* + *α*_0_). We benchmarked the power and false positive rate for DiffDriver and other tests currently used in the field (the same as used in the null simulation section.) The power of DiffDriver was consistently higher than other methods across different simulation settings and sample sizes, while the false positive rate was well under control (**Fig. 3**). In the “low power” setting, where the probability of under selection for a sample is 4-fold in one context than the other, DiffDriver (19.8%) doubles the power of Fisher’s exact test (10.5%) to detect differential selection of a TSG at sample size 1500 (**Fig. 3a**). In the high-power setting, where the probability of under selection for a sample is 9-fold in one context than the other, DiffDriver (39.4%) increases the power by 42% compared to Fisher’s exact test (27.8%) at sample size 1500 (**Fig. 3b**). Similar increase in power is shown for an oncogene (**Fig. 3c-d**). We have also performed simulation analyses for a continuously changing context and observed increase in power for all settings (Supplementary Figure 5). For all these settings, Coselens’s power is on the lower side, comparable to Fisher’s exact test. This is likely because Coselens uses a coarser categorization of functional mutation types (two categories, truncating vs. missense), and fits a separate selection parameter for each category, giving a two-degree-of-freedom test that dilutes power when the signal is concentrated in one category. DiffDriver instead weight multiple functional annotations based on parameters from learned from data (β^f^) and collapse them into a single differential-selection coefficient (*α*_1_) — a one-degree-of-freedom test that is more powerful. For a continuous context, Coselens loses further power because the context must be binarized. Our results thus showed that with a statistical model of differential selection, and leveraging functional information of mutations, DiffDriver is considerably more sensitive than existing methods for detecting context-associated selection.

### DiffDriver recapitulate known context-associated driver genes

We subsequently applied DiffDriver to primary cancer somatic mutation data to validate its ability to identify genes subject to context-associated selection. As a proof-of-concept, we analyzed two distinct histological subtypes of non-small cell lung cancer: lung adenocarcinoma (LUAD) and lung squamous cell carcinoma (LUSC). These subtypes arise from different cellular lineages—mucus-producing cells and squamous epithelial cells, respectively—and exhibit divergent biological profiles and clinical trajectories^23^. Utilizing cohorts from The Cancer Genome Atlas (TCGA) for LUAD (n = 530) and LUSC (n = 174), we evaluated differential selection associated with lung cancer subtypes for genes previously identified as significantly mutated in either subtype (24 driver genes were tested in total; see Supplementary Table 1).

Fitting the DiffDriver BMM model to the 704 TCGA lung tumors used in this analysis, the inferred signatures recover canonical mutational processes. Signature 4 is the C>A tobacco-smoking signature (SBS4) and Signature 3 is the C>T-at-CpG clock-like aging signature (SBS1). The per-sample loadings of signatures vary systematically with clinical features. For example, the smoking signature is enriched in ever-smokers (r = 0.25), the aging signature in never-smokers (r = –0.33) and correlated with age (r = 0.12). The resulting per-sample background rates reproduce the observed data at fine resolution— highly correlated at the individual sample-by-context level (r = 0.88) for both LUAD and LUSC (Supplementary Figure 6). This ensures that individual mutational rate confounders are captured and adjusted by DiffDriver. At a False Discovery Rate (FDR) of 0.1 (Benjamini-Hochberg procedure), DiffDriver identified nine genes exhibiting significant differential selection (Supplementary Table 1). Specifically, *BRAF*, *EGFR*, *STK11*, and *KRAS* showed stronger positive selection in LUAD, all belonging to the MAPK/ERK Signaling Pathway. Conversely, *PTEN*, *PIK3CA*, *CDKN2A*, *NFE2L2*, and *DPPA4* were under more intense selection in LUSC. These findings highlight the distinct evolutionary pressures and genetic etiologies underlying these subtypes, aligning with established molecular characterizations of lung tumorigenesis.

Environmental factors have been shown to shape which drivers are selected. We therefore tested whether DiffDriver could recover selection differences associated with tobacco smoking in LUAD. In TCGA LUAD (393 ever-smokers and 68 never-smokers), the canonical never-smoker–associated oncogene *EGFR* had similar mutation frequencies in the two groups (10.9% in ever-versus 14.7% in never-smokers; Fisher’s exact p = 0.42); DiffDriver, in contrast, identified significantly stronger positive selection of *EGFR* in never-smokers (adjusted *p* = 0.037; Supplementary Figure 7). Because never-smoker tumors carry substantially lower background mutation rates, a comparable mutation frequency reflects stronger underlying selection, an effect DiffDriver captured by explicitly modeling the per-sample background rate. Thus DiffDriver recapitulated the well-established enrichment of *EGFR*-driven tumorigenesis among never-smokers. Notably, a recent analysis of NSCLC cohorts used a likelihood-ratio test and found no *EGFR*–smoking association^24^; this mirrors our benchmark, in which a similar test (logistic-regression) applied to the same LUAD data also missed the signal (Supplementary Figure 7), underscoring that explicitly modeling mutation rate can improve power.

Then we asked whether DiffDriver could disentangle differential selection from heterogenous mutational processes. Endometrial carcinoma (UCEC) has a subset of tumors hypermutated owing to *POLE* proofreading deficiency or mismatch-repair loss, elevating mutation burden by orders of magnitude. Testing UCEC driver genes together with *POLE* and *POLD1* for differential selection between hypermutated (>1,000 mutations; n = 74) and low-burden (n = 438) tumors, DiffDriver recovered *POLE*—whose exonuclease-domain mutations cause the ultramutator phenotype—as the strongest hypermutator-associated gene (adjusted *p* = 4×10^-17^), together with *POLD1* and *FAT1*, providing a positive control (Supplementary Figure 8a). Conversely, *TP53*, *KRAS*, *CTNNB1*, and *PIK3CA*—the canonical drivers of the non-hypermutated subtypes—were under stronger selection in low-burden tumors. Critically, genes such as *ARID1A* and *PTEN*, which have far higher raw mutation frequencies in hypermutated tumors (Fisher’s exact *p* = 2×10^-25^ and 4×10^-13^), were not considered as under stronger selection in hypermutated tumors (Supplementary Figure 8b), because DiffDriver attributes their elevated frequencies to the higher background mutation rate rather than to stronger selection. This demonstrates that DiffDriver’s per-sample background model separates true differential selection from mutation-rate heterogeneity.

### DiffDriver identifies patterns of differential selection associated with tumor clinical traits

We next applied DiffDriver to investigate genes subject to differential selection across various clinical and molecular traits within the TCGA pan-cancer cohort. These traits included patient age, disease-specific survival, progression-free survival, tumor mutational burden (TMB), fraction of genome altered (FGA), and aneuploidy score. We note again that DiffDriver only tests for association and not causality, and some of the traits tested (e.g. survival) are more likely to be a consequence of differential selection rather than the cause if we see an association (see Discussion). For each of the 20 cancer types analyzed, we restricted our testing to previously identified driver genes in corresponding tumor type^5^. 24.3% of genes are identified as showing differential selection for at least one of these 6 traits (45 out of 185 genes, significance was assessed at Benjamini–Hochberg procedure FDR < 0.1 correcting across tumor types within each trait). Next, we examined results for each tumor type, at FDR < 0.1 correcting for tests performed in that tumor type. The landscape of differential selection across these traits in each tumor type is summarized in **Fig. 4a**. We observed that the number of significant genes identified was positively correlated with tumor cohort size, suggesting that our power to detect differential selection remains constrained in smaller datasets. Of all genes tested, TP53 exhibited trait-associated selection in the largest number of tumor types. As shown in **Fig. 4b** (left), the selection pressure on *TP53* varied predictably with clinical and genomic features: *TP53* was more frequently inferred to be under selection in tumors with shorter survival and lower TMB, and with higher inferred selection intensity as genomic instability (FGA and aneuploidy scores) increased. These two associations likely have different origins. The positive association with genomic instability is likely causal in the reverse direction: extensive experimental evidence shows that loss of p53 function drives chromosomal instability and aneuploidy^25,26^—through abrogated cell-cycle checkpoints, tolerance of DNA damage, and centrosome amplification, so genome instability is likely a downstream consequence of TP53 inactivation than as a context that selects for it. By contrast, the negative association with TMB likely has a different explanation. Genome instability and TMB have been found to be negatively correlated across tumors^27,28^. A potential explanation is that a tumor can acquire the alterations needed for malignancy through one of two largely alternative routes—an elevated point-mutation rate (high TMB; e.g., from mismatch-repair or polymerase-proofreading defects) or chromosomal instability (high FGA/aneuploidy; e.g., from TP53 loss). Because only one route is generally sufficient to reach malignancy, conditioning on tumor formation—a common downstream effect of both—induces a spurious negative correlation between them (a case of collider bias).

**Fig. 4.**
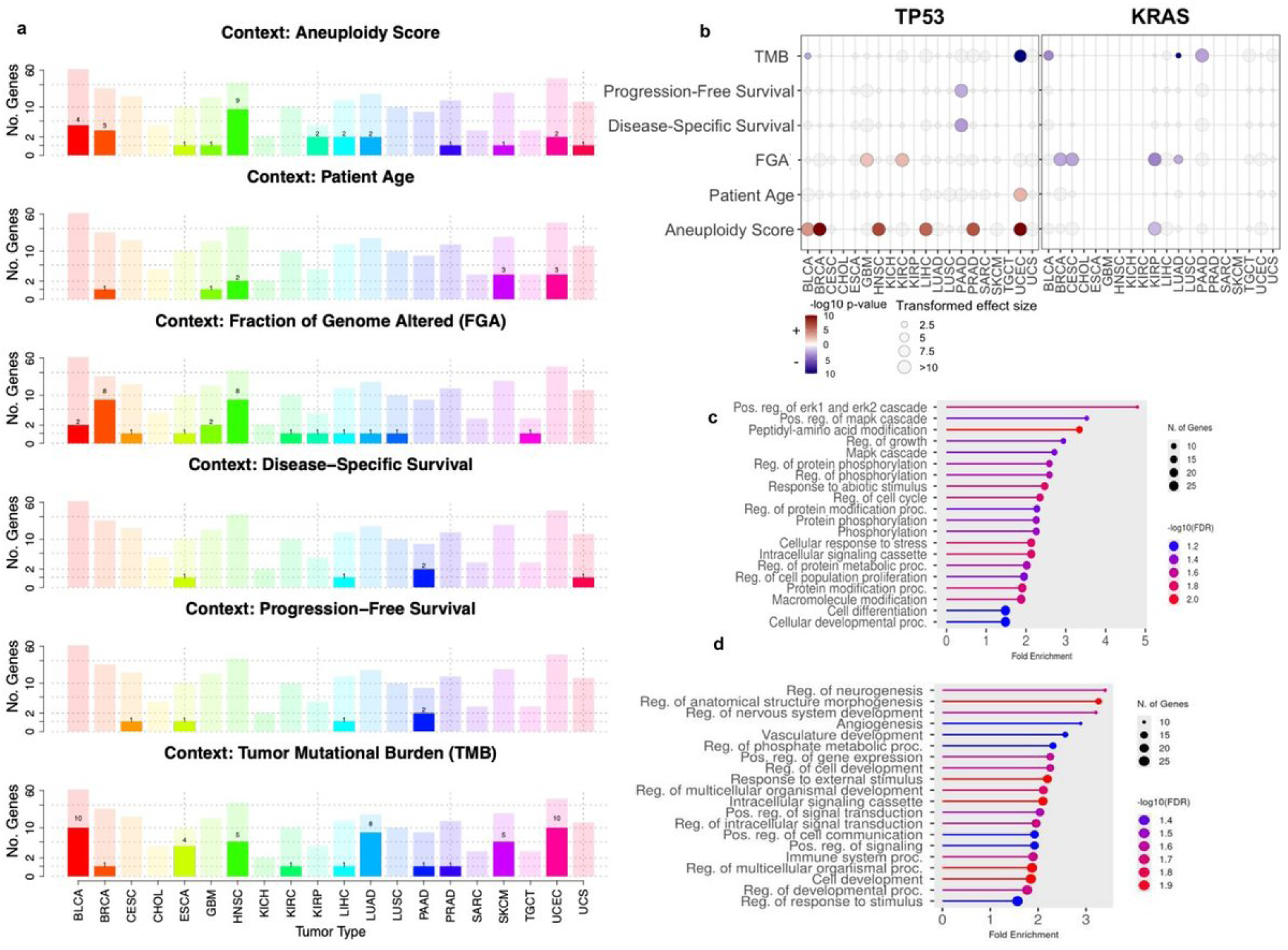
DiffDriver identifies context-associated driver genes and functional associations across cancer types. **a,** Number of driver genes identified by DiffDriver associated with six distinct clinical contexts (Aneuploidy Score, Patient Age, Fraction of Genome Altered [FGA], Disease-Specific Survival, Progression-Free Survival, and Tumor Mutational Burden [TMB]) across TCGA tumor types. Solid bars represent the count of significant context-associated driver genes, overlaid on semi-transparent bars indicating the total number of driver genes identified for each tumor type. **b,** Pan-cancer association landscape for *TP53* and *KRAS*. The bubble plot displays the association of mutations in these genes with various clinical phenotypes across different tumor types. Dot color indicates directional significance (red: positive association/risk; blue: negative association/protective, scaled by signed −log_10_ *P* value), and dot size represents the probability ratio of a sample being under selection at the 75th percentile of the cohort’s context value compared to the 25th percentile (or the 25th versus the 75th for negative effect sizes), obtained based on *α*_1_ estimated by DiffDriver. **c, d,** Gene Ontology (GO) enrichment analysis of the identified context-associated driver genes. The plots highlight significant enrichment in (**c**) oncogenic signaling and metabolic pathways (e.g., Ras signaling, cell cycle regulation) and (**d**) immune-related processes (e.g., cytokine production, immune system processes). The x-axis denotes fold enrichment, dot size indicates the number of genes, and color represents statistical significance (−log_10_ FDR).

In contrast, our analysis of *KRAS* (**Fig. 4b**, right) revealed that the KRAS selection signal is elevated in tumors with lower burden and lower genomic instability. A possible explanation is: as a strong MAPK/ERK-activating and frequently initiating driver, KRAS confers its largest fitness gain in genomically stable tumors, whereas in high-burden, high-instability tumors co-occurring drivers and copy-number alterations provide redundant oncogenic routes that dilute its marginal advantage.^17,29^. Consistent with this, alterations activating the same RTK/RAS/MAPK pathway occupy the high-instability niche and are largely mutually exclusive with KRAS^30^: in TCGA stomach adenocarcinoma, receptor tyrosine kinase amplifications (EGFR/ERBB2/MET; 19% of tumors) co-occur with KRAS mutation in only 2 of 438 cases (Fisher p = 0.008) and arise in significantly more genomically unstable tumors (fraction genome altered 0.36 vs 0.19; aneuploidy score 15.5 vs 8.8)^31,32^. Analogous mutual exclusivity of BRAF with KRAS in hypermutated, microsatellite-unstable colorectal cancer^33^ reflects the same redundancy, though the exact drivers differ by tumor type.

We next sought to identify functional commonalities among genes subject to the identified trait-associated differential selection in multiple tumor types. To achieve this, we performed a gene set enrichment analysis (GSEA) by aggregating unique genes identified as differentially selected in each 20 TCGA cohorts (FDR < 0.1 with the Benjamini–Hochberg correction applied across tumor types within each trait), using the total set of tested driver genes as the background (n = 192 unique genes) (Supplementary Tables 2-3). We focused this analysis on the two traits with the highest number of differential selection cases: Tumor Mutational Burden (TMB) (n = 29 unique genes) and genomic instability (defined as a union of genes associated with FGA and aneuploidy score; n = 30 unique genes). For TMB-associated genes (**Fig. 4c**), Gene Ontology (GO) analysis revealed enrichment in cell-growth and signaling pathways (Ras and MAPK/ERK signaling, growth-factor responses, and cell-cycle regulation). Genes associated with genomic instability were enriched across a broad range of processes— including signal transduction, cell migration, and developmental and metabolic processes—but notably in immune and defense responses (**Fig. 4d**). These genes include canonical immune genes *HLA-A*, *HLA-B* and common driver genes with immune functions.

### DiffDriver identifies genes under differential selection associated with tumor immune microenvironments

The tumor immune microenvironment and a tumor’s driver alterations have profound relationship and has been an important topic^12,13^. They can be linked in two directions—the immune context can select for driver alterations (e.g., those that evade immune detection), and driver mutations can in turn shape the immune microenvironment—and both directions are informative, since identifying which drivers associate with which immune contexts can nominate immune-evasion genes and context-specific dependencies relevant to immunotherapy. Prior pan-cancer studies examined such links using mutation frequency^12,13^; here we instead test for differential selection across the previously defined immune subtypes (C1–C6) using DiffDriver. We applied DiffDriver to investigate differential selection patterns across distinct tumor microenvironments, utilizing previously defined pan-cancer immune subtypes (C1–C6)^12^. We focused on 4 subtypes with the most samples in the tumor types included in this study: C1 (wound healing) is characterized by the elevated expression of angiogenic genes and high cellular proliferation; C2 (IFN-*γ* dominant) exhibits the highest M1/M2 macrophage polarization, a robust CD8 signal, and the highest fraction of tumor-infiltrating lymphocytes (TILs); C3 (inflammatory) is defined by elevated Th17 and Th1 gene signatures; and C4 (lymphocyte depleted) is marked by a prominent macrophage signature and a relative absence of lymphocytic infiltrate.

To characterize immune-associated differential selection at the pan-cancer level, we pooled samples across tumor types and tested each driver gene for differential selection between each immune subtype (C1–C4) and the remaining subtypes, adjusting for patient age, sex, and tumor type. We identified 20 gene–immune-subtype associations at an FDR of 0.1, involving 18 unique genes out of 220 genes tested (8.2%, Fig. 5a, Supplementary Table 5). The IFN-γ–dominant subtype C2 accounted for most associations (11 genes): the antigen-presentation genes *HLA-A* and *HLA-B* were both under stronger positive selection in C2 (adjusted *p* = 0.072 and 0.079), consistent with pressure to disable antigen presentation in this lymphocyte-rich, immunologically active subtype, as were the immune regulators *CASP8* and *JAK2* and the tumor suppressor *TP53*; *KRAS*, by contrast, was more strongly selected outside C2 (adjusted *p* = 0.002). In the inflammatory subtype C3, *TP53*, *SMARCA4*, *BRCA1*, *NSD2*, *FAT1*, *LATS2*, and *POLQ* were preferentially selected in the remaining subtypes, and in the lymphocyte-depleted subtype C4, *EGFR* was selected within the subtype while *CIC* was selected outside it. No gene reached significance for the wound-healing subtype C1.

**Fig. 5.**
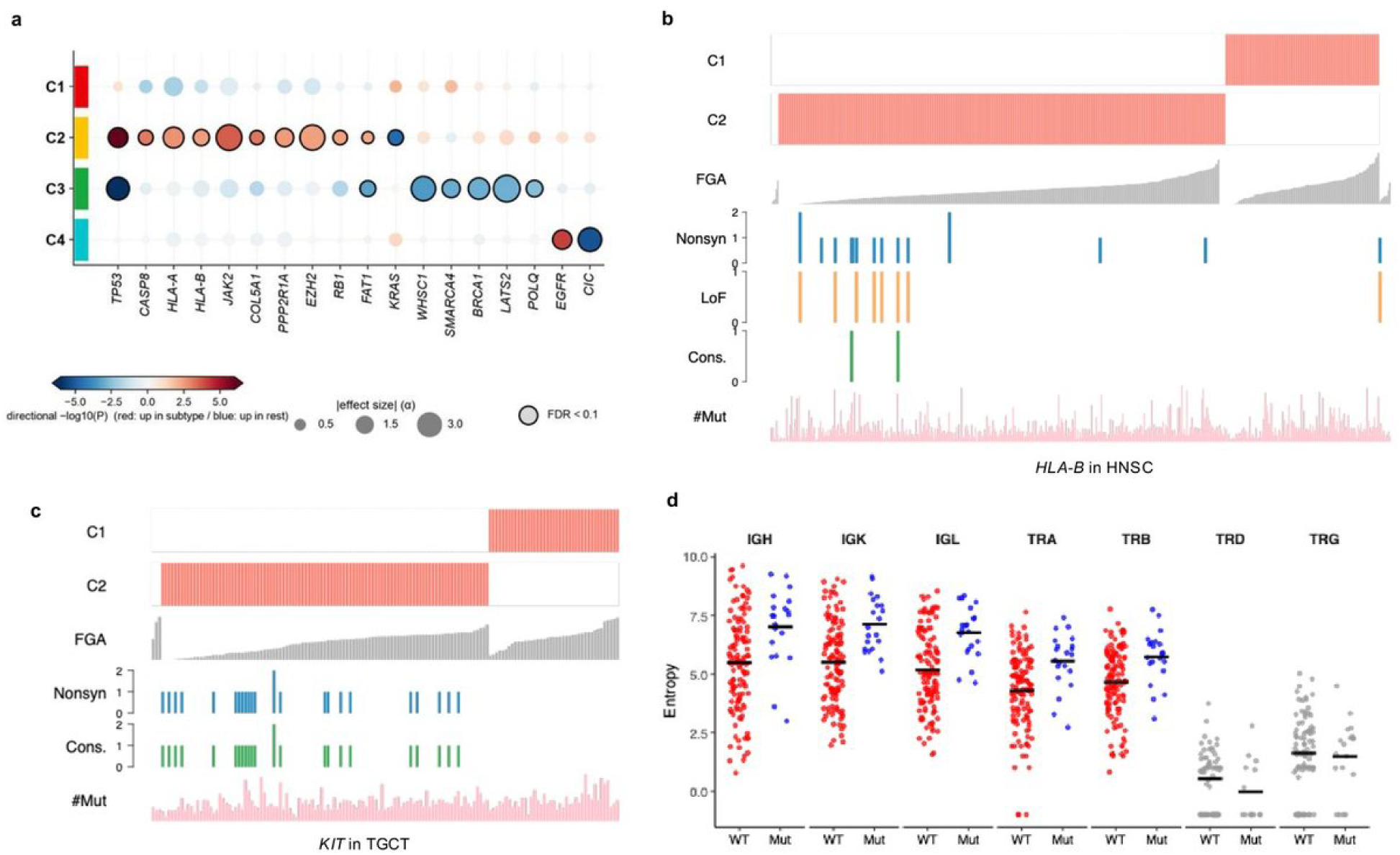
DiffDriver reveals the pan-cancer landscape of driver genes and their association with molecular subtypes and immune heterogeneity. **a,** Pan-cancer landscape of differential selection across immune subtypes. Each dot is a driver gene (columns) tested for differential selection between an immune subtype (rows, C1–C4) and the remaining subtypes, pooling 8,253 tumors across 27 TCGA cancer types. Dot color shows directional significance (−log10 P; red, stronger selection within the subtype; blue, stronger selection in the remaining subtypes), dot size the effect size (|α|, the subtype coefficient), and a black outline marks associations significant at FDR < 0.1; only genes significant in at least one subtype are shown. **b, c,** Detailed visualization of molecular subtypes defined by mutations in *HLA-B* (**b**) and *KIT* (**c**). Top bars indicate cluster assignments (C1, C2). We only displayed C1 and C2, as there are very few samples assigned to other subtypes in this tumor type. Middle tracks display the Fraction Genome Altered (FGA) and specific mutation types (Nonsyn, nonsynonymous; LoF, loss-of-function; Cons., conservative) for each individual. The bottom histograms represent the total mutation burden (#Mut). **d,** Association between *KIT* mutation status and immune repertoire diversity. Dot plots compare the Shannon entropy of various immunoglobulin (Ig) and T-cell receptor (TCR) chains (IGH, IGK, IGL, TRA, TRB, TRD, TRG) between *KIT* wild-type (WT) and mutant (Mut) samples. Horizontal black lines indicate the mean entropy.

We next asked how these pan-cancer signals manifest within individual tumor types (Supplementary Table 4). The antigen-presentation signal was most significant in head and neck squamous cell carcinoma (HNSC), which we examine in detail below (**Fig. 5b**). *HLA-B* is a critical component of the MHC Class I complex; mutations in *HLA-B* can disrupt antigen presentation, thereby facilitating tumor evasion from immune surveillance. Consistently, we found that *HLA-B* is under stronger positive selection within the C2 (IFN -dominant) immune subtype (DiffDriver adjusted *p* value = 0.056). Given that C2 is characterized by a high fraction of tumor-infiltrating lymphocytes and active immune signaling, the selective pressure to disable antigen presentation is likely more in this context compared to others. This interpretation aligns with experimental and genomic evidence that antigen-presentation genes are preferentially disrupted under immune pressure: mutations in HLA and B2M are enriched in tumors with high cytolytic activity^13^, allele-specific HLA loss is a recurrent immune-escape event in the evolution of immune-infiltrated lung tumors^34^, and loss of MHC class I antigen presentation (e.g., via B2M) can confer acquired resistance to PD-1 checkpoint blockade^35^.

Beyond canonical immune genes, DiffDriver identified genes with less-characterized roles in the tumor-immune interface, such as the receptor tyrosine kinase *KIT* in tumor type Testicular Germ Cell Tumors (TGCT). Although typically recognized as an oncogene promoting proliferation, we found *KIT* under stronger positive selection within the C2 subtype (DiffDriver adjusted *p* value = 5.22 × 10^−5^, **Fig. 5c**). We further characterized immune repertoire diversity by calculating the entropy for B-cell (IGH, IGK, IGL) and T-cell (TRA, TRB, TRD, TRG) receptors. We observed that higher entropy values—indicating a more diverse and polyclonal immune landscape with a broader array of distinct clonotypes—were positively associated with *KIT* selection strength. This suggests that intensified evolutionary pressure on the *KIT* driver gene is accompanied by a more active and varied adaptive immune response within the tumor microenvironment (**Fig. 5d**). These results suggest *KIT* may facilitate immune evasion or modulate the tumor microenvironment. Notably, *KRAS* and *NRAS* in TGCT were also under stronger selection in the C2 subtype. *KIT* and *RAS* mutations were largely mutually exclusive but collectively affected 30% of TGCT samples (Supplementary Figure 9, suggesting *KIT* may play a functional role similar to *RAS* in TGCT pathogenesis and immune evasion. This immune-modulatory role is experimentally supported in another tumor type: gastrointestinal stromal tumor (GIST). In GIST, oncogenic *KIT* signaling promotes immune escape by upregulating IDO and sustaining regulatory T cells, and *KIT* inhibition with imatinib activates intratumoral CD8+ T cells while inducing regulatory T-cell apoptosis^36^. Concordantly, *KIT*-mutant GISTs display markedly lower cytolytic activity and immune-cell infiltration than *PDGFRA*-mutant tumors^37^, indicating that *KIT* genotype shapes the tumor immune microenvironment. These data may provide a mechanistic basis for the stronger selection of *KIT* within the immune-active C2 subtype that DiffDriver detects in TGCT.

## Discussion

We present a model-based framework to detect differential selection of driver genes across diverse biological contexts. By modeling selection strength as a function of individual-level factors, DiffDriver distinguishes itself from existing selection models that often neglect inter-individual heterogeneity. Through simulations, we demonstrated that the integration of functional annotations significantly improves statistical power. Furthermore, the model effectively accounts for unknown confounders by learning individual-and position-specific background mutation rates.

Notably, DiffDriver identifies associations but cannot inherently determine causality, a limitation shared by all standard association tests. In practice, DiffDriver can evaluate the association between any individual-level covariate and the selection intensity of a driver gene. Such associations may arise from a pre-existing context exerting differential selective pressure (e.g., a highly active immune microenvironment selecting for *HLA* mutations). Conversely, they may reflect the phenotypic consequences of the mutations themselves (e.g., *TP53* mutations driving genomic instability, which subsequently triggers immune surveillance). Thus, the direction of effect is uncertain from the association test alone. The association may also result from the correlation of the individual-level factor under test with the causal factor for differential selection. In this case, we may find proxies or biomarkers for the causal factors that induce differential selection. Therefore, interpreting these associations requires consideration of several plausible hypotheses. External information regarding the direction of effect would be helpful to gain further insight into the DiffDriver results. For instance, using germline genetic background (e.g., polygenic risk scores) as the context ensures that the direction of effect must proceed from context to selection, as somatic events cannot alter the germline. Finally, DiffDriver currently uses binary mutation calls; incorporating additional variant-level information—variant allele frequency, read depth, and clonality—into the per-position likelihood could further refine the estimation of selection strength, and this could be a direction for future work.

Our pan-cancer analysis nominated 18 driver genes whose selection strength varies across immune subtypes (**Fig. 5a**). These pan-cancer associations are broadly concordant with the immune-subtype driver map of Thorsson et al.^12^. These two analyses have several differences so we did not expect identical results. Thorsson et al. asked whether a gene was mutated more often in a subtype, whereas DiffDriver asks whether the strength of selection on a gene differs between subtypes. Thorsson et al included more tumor types, samples and tested more driver genes than in this study. 13 of our 18 genes are reported by Thorsson et al., including several genes involved in antigen-presentation and immune system, *HLA-A*, *HLA-B*, and *CASP8*, and other driver genes *TP53*, *KRAS*, *EGFR*, *RB1*, and *SMARCA4*. Five genes are unique to our selection-based analysis (*COL5A1*, *EZH2*, *LATS2*, *NSD2*, and *POLQ*). Several of these have plausible links to tumor immunity: *EZH2* represses MHC-I antigen presentation and interferon-response chemokines^38,39^, *LATS2* loss activates immunosuppressive YAP/TAZ signaling^40^; the extracellular-matrix collagen *COL5A1* contributes to the stromal barrier that restricts T-cell infiltration^41^; and *NSD2* and *POLQ* act in histone-methylation and DNA-damage (microhomology-mediated end-joining) repair, respectively^42,43^. The results associating these 18 genes and the tumor immune microenvironment together highlighted a few themes. One clear pattern was the preferential selection of genes central to immune recognition in the IFN-γ– dominant subtype C2. These include the antigen-presentation genes *HLA-A* and *HLA-B*, the death-receptor adaptor *CASP8*, a key effector of immune-mediated tumor killing^13,44^, and *JAK2*, a central transducer of IFN-γ/JAK–STAT signaling^45^. The selection of antigen-presentation and IFN-signaling genes in the most immunologically active, lymphocyte-rich subtype suggested selective pressure to evade an active anti-tumor immune response. A second group comprises chromatin and epigenetic regulators—*EZH2*, *SMARCA4*, and *NSD2*—increasingly recognized to shape tumor immunogenicity: *SMARCA4* loss reshapes tumor immunogenicity and cytotoxic T-cell infiltration through interferon-pathway activation^46^, and *EZH2* can repress antigen-presentation and immune-response genes^38^. Classic drivers with pleiotropic effects on tumor immunity—*TP53*^47^, *KRAS*, and *EGFR*—were also nominated, together with DNA-damage-response genes (*BRCA1*, *POLQ*). Their associations with the immune microenvironment can be direct or indirect.

Examining individual tumor types sharpened several of these signals. *HLA-B* was most conspicuously selected in the C2 subtype of head and neck squamous cell carcinoma (Fig. 5b), where disruption of antigen presentation would likely be especially advantageous. The receptor tyrosine kinase *KIT* (Fig. 5c–d), though best known as a proliferative oncogene, was under stronger selection in the immune-active C2 subtype of TGCT. Direct experimental evidence has linked *KIT* signaling to remodeling of the tumor immune microenvironment and T-cell evasion in gastrointestinal stromal tumor^36,37^. Together, these results indicate that many cancer driver genes are under immune-context-associated selection, and that DiffDriver nominates candidate mediators of immune evasion, including both canonical immune genes and less-expected driver genes.

Applying DiffDriver to the TCGA cohort, we observed that the number of identified genes increased with sample size, suggesting that we have not yet reached saturation and that larger cohorts are in need. We anticipate broader applications of this tool as datasets of tumor genetic profiles, clinical records, and molecular phenotypes continue to expand. For example, when comparing tumors across different treatment regimens, identifying differential selection can pinpoint key genes driving therapeutic response or the emergence of drug resistance. Ultimately, we anticipate that DiffDriver will serve as a useful tool for characterizing the causes and consequences of inter-individual tumor heterogeneity.

## Data availability

Phenotype data, mutation data, and gene list data are available via Zenodo at https://doi.org/10.5281/zenodo.18521355, https://doi.org/10.5281/zenodo.18521782, and https://doi.org/10.5281/zenodo.18521873, respectively. The package also relies on 9-annotation and 96-annotation files, which are available at https://doi.org/10.5281/zenodo.18462105.

## Code availability

Our software is available at https://github.com/szhaolab/diffdriver. Code related to analyses performed in this study can be accessed at https://github.com/szhaolab/DiffDriver-PAPER.

## Acknowledgement.

We would like to thank Matthew Stephens, Thomas Gajewski, Lixing Yang at University of Chicago, Brock Christensen at Dartmouth College for the discussion of the project. The work is supported by National Institute of Health grants P20GM130454, R35GM154925, an American Cancer Society Institutional Research Grant at Dartmouth to S.Z., and an NIH grant R01AI175554 to X.H..

