## Supplementary Information for "DiffDriver Detects Context-Associated Selection of Cancer Driver Genes"

- an EM algorithm[1] to estimate parameters $\theta = \{\alpha_1, \alpha_0, \beta_0^f\}$

The complete log likelihood is given by

$$\log P(y, Z|\theta) = \sum_i I\,(Z_i = 1)(\log\pi_i + \log P(y_i|\beta^f, \beta_0^f, \mu, x^f, Z_i = 1)) + \sum_i I\,(Z_i = 0)\big(\log(1-\pi_i) + \log P(y_i|\mu, Z_i = 0)\big). \quad (11)$$

The expected value of complete log likelihood can be written as

$$E_{Z|y}\log P(y, Z|\theta) = \sum_i p_i\,(\log\pi_i + \log P(y_i|\beta^f, \beta_0^f, \mu, x^f, Z_i = 1)) + \sum_i (1-p_i)(\log(1-\pi_i) + \log P(y_i|\mu, Z_i = 0)\}. \quad (12)$$

Where $p_i = P(Z_i = 1|y)$ is the posterior probability of sample $i$ is under selection.

- In the E-step, we get the Q function

$$Q(\theta|\theta^{(t)}) = E_{Z|y}\log P(y, Z|\theta^{(t)}). \quad (13)$$

Given the current parameter estimates $\theta^{(t)}$ in the $t$-th iteration, replace the missing data $Z_i$ by the estimated a-posteriori probabilities, $\widehat{p_i} = P(Z_i = 1|y, \theta^{(t)})$.

- In the M-step, we found $\theta$ that maximizes the Q function.

The Q function can be written as

$$Q(\theta|\theta^{(t)}) = Q_1(\alpha, \alpha_0|\theta^{(t)}) + Q_2(\beta_0^f|\theta^{(t)}), \quad (14)$$

where

$$Q_1(\alpha, \alpha_0|\theta^{(t)}) = \sum_i \widehat{p_i}\log\pi_i + \sum_i (1-\widehat{p_i})\log(1-\pi_i), \quad (15)$$

$$Q_2(\beta_0^f|\theta^{(t)}) = \sum_i \widehat{p_i}\log P(y_i|\beta_0, \mu, x^f, Z_i = 1) + \sum_i (1-\widehat{p_i})\log P(y_i|\mu, Z_i = 0). \quad (16)$$

$Q_1$ and $Q_2$ can be maximized separately at each iteration. When maximizing $Q_1$, we fit a firth logistic regression model to instead of a standard logistic regression model, given the mutation data used here is very sparse. To maximize $Q2$, we use the Nelder-Mead method implemented in the optim() function in R.

**Supplementary Table 1 | Differential driver gene analysis between lung adenocarcinoma and squamous cell carcinoma.** DiffDriver was applied to identify driver genes specifically associated with TCGA Lung Adenocarcinoma (LUAD, target phenotype E1) versus Lung Squamous Cell Carcinoma (LUSC, reference phenotype E0). **α**, estimated effect size from the DiffDriver model (positive values indicate a positive association with the LUAD subtype); ***p* value**, nominal *P* value derived from the DiffDriver test; **FDR**, false discovery rate-adjusted *P* value (*q*-value); **# mut. in E = 1**, total count of observed mutations for the specific gene in the LUAD cohort; **# mut. in E = 0**, total count of observed mutations for the specific gene in the LUSC cohort. Genes with FDR < 0.1 are shown in bold. Total sample sizes were fixed at $\mathbf{n_{E1} = 530}$ (LUAD) and $\mathbf{n_{E0} = 174}$ (LUSC).

| Gene | α | *p* value | FDR | # Mut. in E = 1 | # Mut. in E = 0 |
|---|---|---|---|---|---|
| ***KRAS*** | **4.68** | **1.29E-17** | **3.09E-16** | **152** | **2** |
| ***NFE2L2*** | **-3.17** | **9.02E-09** | **1.08E-07** | **15** | **27** |
| ***CDKN2A*** | **-3.08** | **1.05E-05** | **8.39E-05** | **17** | **21** |
| ***STK11*** | **3.26** | **2.45E-05** | **1.47E-04** | **38** | **0** |
| ***DPPA4*** | **-4.81** | **1.24E-04** | **5.97E-04** | **4** | **10** |
| ***PTEN*** | **-2.8** | **5.59E-04** | **2.23E-03** | **8** | **13** |
| ***PIK3CA*** | **-1.6** | **6.49E-04** | **2.23E-03** | **27** | **28** |
| ***EGFR*** | **1.58** | **2.32E-03** | **6.96E-03** | **52** | **5** |
| ***BRAF*** | **1.43** | **1.94E-02** | **5.18E-02** | **36** | **8** |
| *LCE1F* | 2.74 | 5.02E-02 | 1.20E-01 | 11 | 0 |
| *MGA* | 2.14 | 6.43E-02 | 1.31E-01 | 35 | 6 |
| *DZIP1L* | 2.97 | 6.56E-02 | 1.31E-01 | 14 | 0 |
| *SMARCA4* | 1.56 | 7.79E-02 | 1.44E-01 | 40 | 6 |
| *KCNQ2* | 2.29 | 1.01E-01 | 1.73E-01 | 22 | 4 |
| *SETD2* | 0.98 | 1.52E-01 | 2.43E-01 | 32 | 5 |
| *ELTD1* | -0.97 | 2.54E-01 | 3.82E-01 | 34 | 14 |

| | | | | | |
|---|---|---|---|---|---|
| *ARID1A* | 0.92 | 2.85E-01 | 4.03E-01 | 30 | 6 |
| *ARID2* | 1.55 | 3.18E-01 | 4.24E-01 | 25 | 9 |
| *ADAMTS12* | -0.73 | 3.99E-01 | 5.05E-01 | 105 | 29 |
| *KEAP1* | 0.95 | 5.05E-01 | 6.06E-01 | 80 | 21 |
| *NF1* | 0.06 | 1.00E+00 | 1.00E+00 | 47 | 17 |
| *PTPRU* | 0.02 | 1.00E+00 | 1.00E+00 | 16 | 6 |
| *RB1* | -0.81 | 1.00E+00 | 1.00E+00 | 19 | 9 |
| *TP53* | -0.44 | 1.00E+00 | 1.00E+00 | 249 | 128 |

**Supplementary Table 2 | Driver genes associated with tumor mutation burden across 20 TCGA cancer types.** DiffDriver was applied to identify driver genes significantly associated with Tumor Mutation Burden (TMB) across 20 TCGA tumor types. **α**, estimated effect size from the DiffDriver model; ***p* value**, nominal *P* value derived from the DiffDriver test; **FDR**, false discovery rate-adjusted *P* value (*q*-value); **Tumor**, the specific TCGA cancer type where the association was identified. Tumor mutational burden (TMB) is used as the genomic phenotype. Only genes with FDR < 0.1 are included.

| Gene | α | *p* value | FDR | # Mut. in E = 1 | # Mut. in E = 0 | # E = 1 | # E = 0 | Tumor | FDR (per-trait BH) |
|---|---|---|---|---|---|---|---|---|---|
| *FGFR3* | -0.24 | 8.29E-10 | 5.22E-08 | 16 | 44 | 133 | 258 | BLCA | 4.53E-08 |
| *NFE2L2* | -0.31 | 3.52E-05 | 1.11E-03 | 5 | 19 | 133 | 258 | BLCA | 9.62E-04 |
| *KRAS* | -0.37 | 1.68E-04 | 3.54E-03 | 2 | 11 | 133 | 258 | BLCA | 4.09E-03 |
| *HRAS* | -0.22 | 9.12E-04 | 1.37E-02 | 3 | 12 | 133 | 258 | BLCA | 1.49E-02 |
| *CREBBP* | -0.25 | 1.09E-03 | 1.37E-02 | 16 | 25 | 133 | 258 | BLCA | 1.70E-02 |
| *TP53* | -0.18 | 1.71E-03 | 1.79E-02 | 91 | 106 | 133 | 258 | BLCA | 2.43E-02 |
| *TSC1* | -0.22 | 4.71E-03 | 4.24E-02 | 12 | 16 | 133 | 258 | BLCA | 5.36E-02 |
| *FBXW7* | -0.12 | 1.17E-02 | 9.04E-02 | 14 | 16 | 133 | 258 | BLCA | 9.86E-02 |
| *RHOB* | -0.11 | 1.29E-02 | 9.04E-02 | 6 | 23 | 133 | 258 | BLCA | 1.03E-01 |
| *CDKN1A* | -0.26 | 1.52E-02 | 9.59E-02 | 4 | 9 | 133 | 258 | BLCA | 1.16E-01 |
| *MAP3K1* | -0.82 | 3.94E-04 | 9.84E-03 | 2 | 24 | 267 | 652 | BRCA | 8.07E-03 |
| *NFE2L2* | -0.90 | 5.72E-04 | 5.72E-03 | 1 | 15 | 60 | 120 | ESCA | 1.04E-02 |

| | | | | | | | | | |
|---|---|---|---|---|---|---|---|---|---|
| *CDKN2A* | -1.52 | 1.79E-03 | 8.93E-03 | 0 | 10 | 60 | 120 | ESCA | 2.44E-02 |
| *TGFBR2* | -0.76 | 1.32E-02 | 4.41E-02 | 2 | 6 | 60 | 120 | ESCA | 1.03E-01 |
| *SLC17A6* | -0.72 | 2.18E-02 | 5.46E-02 | 1 | 7 | 60 | 120 | ESCA | 1.56E-01 |
| *CDKN2A* | -0.33 | 1.52E-05 | 5.01E-04 | 24 | 56 | 158 | 333 | HNSC | 4.78E-04 |
| *NOTCH1* | -0.24 | 3.38E-04 | 5.29E-03 | 25 | 44 | 158 | 333 | HNSC | 7.39E-03 |
| *PIK3CA* | -0.19 | 4.80E-04 | 5.29E-03 | 29 | 57 | 158 | 333 | HNSC | 9.27E-03 |
| *NSD1* | 0.22 | 9.77E-03 | 7.74E-02 | 34 | 13 | 158 | 333 | HNSC | 8.90E-02 |
| *FBXW7* | -0.30 | 1.17E-02 | 7.74E-02 | 8 | 16 | 158 | 333 | HNSC | 9.86E-02 |
| *PBRM1* | -2.33 | 8.62E-03 | 8.62E-02 | 35 | 28 | 166 | 162 | KIRC | 8.57E-02 |
| *KEAP1* | 0.68 | 2.25E-03 | 3.15E-02 | 11 | 3 | 143 | 210 | LIHC | 2.95E-02 |
| *EGFR* | -0.56 | 2.65E-15 | 5.03E-14 | 13 | 36 | 186 | 339 | LUAD | 4.34E-13 |
| *KRAS* | -0.14 | 8.35E-14 | 7.93E-13 | 56 | 96 | 186 | 339 | LUAD | 6.85E-12 |
| *STK11* | -0.17 | 3.66E-06 | 2.32E-05 | 18 | 20 | 186 | 339 | LUAD | 1.36E-04 |
| *KEAP1* | -0.11 | 1.60E-05 | 7.61E-05 | 30 | 50 | 186 | 339 | LUAD | 4.78E-04 |
| *SETD2* | -0.10 | 4.01E-03 | 1.52E-02 | 16 | 16 | 186 | 339 | LUAD | 4.87E-02 |
| *BRAF* | -0.06 | 8.21E-03 | 2.60E-02 | 20 | 16 | 186 | 339 | LUAD | 8.41E-02 |
| *NFE2L2* | -0.10 | 2.33E-02 | 5.59E-02 | 7 | 8 | 186 | 339 | LUAD | 1.57E-01 |
| *RB1* | -0.11 | 2.35E-02 | 5.59E-02 | 10 | 9 | 186 | 339 | LUAD | 1.57E-01 |

| | | | | | | | | | |
|---|---|---|---|---|---|---|---|---|---|
| *KRAS* | -2.82 | 1.31E-03 | 1.05E-02 | 42 | 43 | 51 | 69 | PAAD | 1.95E-02 |
| *CDK12* | 2.14 | 6.65E-03 | 9.31E-02 | 7 | 3 | 179 | 304 | PRAD | 7.04E-02 |
| *NRAS* | -0.08 | 3.73E-06 | 7.46E-05 | 33 | 56 | 99 | 182 | SKCM | 1.36E-04 |
| *BRAF* | -0.03 | 3.06E-03 | 3.06E-02 | 58 | 100 | 99 | 182 | SKCM | 3.86E-02 |
| *PTEN* | -0.07 | 4.74E-03 | 3.16E-02 | 7 | 9 | 99 | 182 | SKCM | 5.36E-02 |
| *NF1* | 0.11 | 2.35E-02 | 1.00E-01 | 28 | 8 | 99 | 182 | SKCM | 1.57E-01 |
| *PPP6C* | -0.05 | 2.50E-02 | 1.00E-01 | 6 | 12 | 99 | 182 | SKCM | 1.63E-01 |
| *TP53* | -0.49 | 6.70E-16 | 2.68E-14 | 9 | 44 | 71 | 153 | UCEC | 2.20E-13 |
| *CTNNB1* | -0.26 | 1.01E-14 | 2.01E-13 | 12 | 56 | 71 | 153 | UCEC | 1.10E-12 |
| *CHD4* | -0.93 | 5.34E-10 | 7.12E-09 | 5 | 19 | 71 | 153 | UCEC | 3.50E-08 |
| *PPP2R1A* | -0.29 | 3.37E-06 | 3.37E-05 | 5 | 15 | 71 | 153 | UCEC | 1.36E-04 |
| *ARID1A* | -0.20 | 1.74E-04 | 1.40E-03 | 19 | 25 | 71 | 153 | UCEC | 4.09E-03 |
| *CTCF* | -0.14 | 7.51E-04 | 5.01E-03 | 11 | 17 | 71 | 153 | UCEC | 1.30E-02 |
| *FGFR2* | -0.08 | 6.50E-03 | 3.72E-02 | 11 | 10 | 71 | 153 | UCEC | 7.04E-02 |
| *NRAS* | -0.20 | 8.96E-03 | 4.23E-02 | 4 | 2 | 71 | 153 | UCEC | 8.64E-02 |
| *PIK3CA* | -0.06 | 9.52E-03 | 4.23E-02 | 53 | 82 | 71 | 153 | UCEC | 8.90E-02 |
| *SPOP* | -0.12 | 1.10E-02 | 4.41E-02 | 4 | 12 | 71 | 153 | UCEC | 9.77E-02 |

**Supplementary Table 3 | Driver genes associated with genome instability across 20 TCGA cancer types.** DiffDriver was applied to identify driver genes significantly associated with genome instability metrics, specifically Aneuploidy Score and Fraction Genome Altered (FGA), across 20 TCGA tumor types. **α**, estimated effect size from the DiffDriver model; ***p* value**, nominal *P* value derived from the DiffDriver test; **FDR**, false discovery rate-adjusted *P* value (*q*-value); **Tumor**, the specific TCGA cancer type where the association was identified; **Context**, genomic phenotype (Aneuploidy Score or Fraction Genome Altered). Only genes with FDR < 0.1 are included.

| Gene | α | *p* value | FDR | # Mut. in E = 1 | # Mut. in E = 0 | # E = 1 | # E = 0 | Tumor | Context | FDR (per-trait BH) |
|---|---|---|---|---|---|---|---|---|---|---|
| *FGFR3* | -0.12 | 6.68E-07 | 4.21E-05 | 20 | 38 | 208 | 173 | BLCA | Aneuploidy Score | 3.24E-05 |
| *HRAS* | -0.18 | 8.02E-05 | 2.53E-03 | 5 | 10 | 208 | 173 | BLCA | Aneuploidy Score | 2.15E-03 |
| *TP53* | 0.52 | 1.80E-04 | 3.79E-03 | 130 | 65 | 208 | 173 | BLCA | Aneuploidy Score | 4.22E-03 |
| *NFE2L2* | -0.13 | 7.14E-04 | 1.12E-02 | 11 | 13 | 208 | 173 | BLCA | Aneuploidy Score | 1.30E-02 |
| *TP53* | 0.68 | 7.93E-13 | 1.98E-11 | 150 | 71 | 405 | 492 | BRCA | Aneuploidy Score | 1.30E-10 |
| *CTCF* | -0.43 | 6.92E-07 | 8.66E-06 | 0 | 14 | 405 | 492 | BRCA | Aneuploidy Score | 3.24E-05 |
| *MAP3K1* | -0.11 | 5.27E-03 | 4.39E-02 | 7 | 19 | 405 | 492 | BRCA | Aneuploidy Score | 7.52E-02 |
| *SMARCA4* | -0.25 | 5.70E-03 | 5.70E-02 | 2 | 8 | 81 | 79 | ESCA | Aneuploidy Score | 7.79E-02 |
| *IDH1* | -0.69 | 4.62E-04 | 7.39E-03 | 2 | 11 | 73 | 189 | GBM | Aneuploidy Score | 9.46E-03 |
| *HRAS* | -0.29 | 6.42E-11 | 2.12E-09 | 2 | 31 | 214 | 268 | HNSC | Aneuploidy Score | 7.02E-09 |
| *TP53* | 0.70 | 1.05E-07 | 1.73E-06 | 172 | 158 | 214 | 268 | HNSC | Aneuploidy Score | 8.57E-06 |
| *CASP8* | -0.18 | 3.50E-06 | 3.14E-05 | 7 | 36 | 214 | 268 | HNSC | Aneuploidy Score | 1.25E-04 |
| *RAC1* | -0.45 | 3.80E-06 | 3.14E-05 | 1 | 12 | 214 | 268 | HNSC | Aneuploidy Score | 1.25E-04 |

| | | | | | | | | | | |
|---|---|---|---|---|---|---|---|---|---|---|
| *HLA-B* | -0.54 | 4.35E-05 | 2.87E-04 | 2 | 14 | 214 | 268 | HNSC | Aneuploidy Score | 1.30E-03 |
| *TGFBR2* | -0.16 | 1.02E-03 | 5.62E-03 | 5 | 17 | 214 | 268 | HNSC | Aneuploidy Score | 1.68E-02 |
| *CTCF* | -0.18 | 1.28E-02 | 6.03E-02 | 1 | 11 | 214 | 268 | HNSC | Aneuploidy Score | 1.50E-01 |
| *EP300* | -0.13 | 2.29E-02 | 8.67E-02 | 9 | 23 | 214 | 268 | HNSC | Aneuploidy Score | 2.21E-01 |
| *HLA-A* | -0.13 | 2.36E-02 | 8.67E-02 | 3 | 11 | 214 | 268 | HNSC | Aneuploidy Score | 2.21E-01 |
| *KRAS* | -0.33 | 7.13E-03 | 2.85E-02 | 0 | 5 | 116 | 158 | KIRP | Aneuploidy Score | 9.35E-02 |
| *CUL3* | -0.39 | 1.69E-02 | 3.38E-02 | 1 | 8 | 116 | 158 | KIRP | Aneuploidy Score | 1.85E-01 |
| *TP53* | 1.14 | 2.27E-06 | 3.17E-05 | 46 | 40 | 130 | 218 | LIHC | Aneuploidy Score | 9.30E-05 |
| *HNF1A* | -0.56 | 7.89E-04 | 5.52E-03 | 1 | 8 | 130 | 218 | LIHC | Aneuploidy Score | 1.36E-02 |
| *SETD2* | -0.15 | 4.11E-04 | 4.73E-03 | 9 | 20 | 242 | 220 | LUAD | Aneuploidy Score | 8.99E-03 |
| *BRAF* | -0.13 | 4.97E-04 | 4.73E-03 | 16 | 20 | 242 | 220 | LUAD | Aneuploidy Score | 9.60E-03 |
| *TP53* | 0.80 | 3.81E-07 | 5.33E-06 | 30 | 12 | 130 | 331 | PRAD | Aneuploidy Score | 2.50E-05 |
| *MAP2K1* | 0.13 | 4.83E-03 | 9.66E-02 | 13 | 3 | 113 | 164 | SKCM | Aneuploidy Score | 7.20E-02 |
| *TP53* | 0.39 | 1.28E-18 | 5.14E-17 | 36 | 16 | 69 | 147 | UCEC | Aneuploidy Score | 4.21E-16 |
| *PPP2R1A* | 0.17 | 8.53E-05 | 1.71E-03 | 12 | 6 | 69 | 147 | UCEC | Aneuploidy Score | 2.15E-03 |
| *PTEN* | -0.24 | 4.66E-03 | 6.06E-02 | 2 | 8 | 31 | 22 | UCS | Aneuploidy Score | 7.20E-02 |
| *FGFR3* | -3.73 | 1.80E-04 | 9.52E-03 | 21 | 39 | 178 | 208 | BLCA | FGA | 5.90E-03 |
| *HRAS* | -6.69 | 3.02E-04 | 9.52E-03 | 3 | 12 | 178 | 208 | BLCA | FGA | 9.01E-03 |

| | | | | | | | | | | |
|---|---|---|---|---|---|---|---|---|---|---|
| *PIK3CA* | -5.13 | 2.78E-10 | 6.94E-09 | 109 | 208 | 417 | 502 | BRCA | FGA | 3.50E-08 |
| *CDH1* | -6.94 | 6.23E-06 | 7.78E-05 | 15 | 35 | 417 | 502 | BRCA | FGA | 4.08E-04 |
| *CTCF* | -11.57 | 1.76E-05 | 1.47E-04 | 1 | 13 | 417 | 502 | BRCA | FGA | 9.62E-04 |
| *MAP3K1* | -6.57 | 3.86E-05 | 2.41E-04 | 6 | 20 | 417 | 502 | BRCA | FGA | 1.81E-03 |
| *KRAS* | -13.50 | 2.16E-03 | 1.08E-02 | 0 | 5 | 417 | 502 | BRCA | FGA | 3.75E-02 |
| *CBFB* | -6.51 | 5.16E-03 | 2.15E-02 | 3 | 9 | 417 | 502 | BRCA | FGA | 7.05E-02 |
| *RUNX1* | -6.89 | 7.65E-03 | 2.73E-02 | 3 | 9 | 417 | 502 | BRCA | FGA | 9.54E-02 |
| *MAP2K4* | -5.36 | 1.49E-02 | 4.66E-02 | 6 | 13 | 417 | 502 | BRCA | FGA | 1.53E-01 |
| *KRAS* | -9.91 | 1.49E-03 | 2.52E-02 | 1 | 10 | 68 | 112 | CESC | FGA | 3.33E-02 |
| *NFE2L2* | 7.84 | 2.22E-03 | 2.22E-02 | 11 | 5 | 89 | 91 | ESCA | FGA | 3.75E-02 |
| *NF1* | -20.18 | 7.28E-03 | 7.15E-02 | 3 | 16 | 96 | 173 | GBM | FGA | 9.54E-02 |
| *TP53* | 7.66 | 8.94E-03 | 7.15E-02 | 43 | 41 | 96 | 173 | GBM | FGA | 1.05E-01 |
| *HRAS* | -18.91 | 1.71E-14 | 5.63E-13 | 0 | 33 | 218 | 269 | HNSC | FGA | 5.59E-12 |
| *CASP8* | -14.12 | 3.20E-10 | 5.28E-09 | 5 | 39 | 218 | 269 | HNSC | FGA | 3.50E-08 |
| *RAC1* | -37.92 | 2.53E-06 | 2.78E-05 | 1 | 12 | 218 | 269 | HNSC | FGA | 2.08E-04 |
| *HLA-B* | -20.68 | 7.51E-05 | 6.20E-04 | 2 | 14 | 218 | 269 | HNSC | FGA | 3.08E-03 |
| *EPHA2* | -25.56 | 6.08E-04 | 4.01E-03 | 0 | 13 | 218 | 269 | HNSC | FGA | 1.66E-02 |
| *NOTCH1* | -4.94 | 1.60E-03 | 7.65E-03 | 26 | 42 | 218 | 269 | HNSC | FGA | 3.33E-02 |

| | | | | | | | | | | |
|---|---|---|---|---|---|---|---|---|---|---|
| *TGFBR2* | -6.31 | 1.62E-03 | 7.65E-03 | 8 | 14 | 218 | 269 | HNSC | FGA | 3.33E-02 |
| *HLA-A* | -8.25 | 2.22E-03 | 9.17E-03 | 2 | 12 | 218 | 269 | HNSC | FGA | 3.75E-02 |
| *TP53* | 6.57 | 3.94E-03 | 3.94E-02 | 6 | 0 | 139 | 267 | KIRC | FGA | 5.61E-02 |
| *KRAS* | -19.14 | 1.25E-04 | 5.00E-04 | 0 | 5 | 84 | 191 | KIRP | FGA | 4.55E-03 |
| *HNF1A* | -14.53 | 9.71E-04 | 1.36E-02 | 1 | 8 | 145 | 207 | LIHC | FGA | 2.45E-02 |
| *KRAS* | -3.67 | 2.42E-03 | 4.60E-02 | 63 | 78 | 208 | 256 | LUAD | FGA | 3.78E-02 |
| *KEAP1* | 9.32 | 7.85E-03 | 7.85E-02 | 15 | 3 | 80 | 87 | LUSC | FGA | 9.54E-02 |
| *KIT* | -6.89 | 2.29E-03 | 6.86E-03 | 9 | 15 | 86 | 61 | TGCT | FGA | 3.75E-02 |

**Supplementary Table 4 | Significant differential driver genes associated with immune subtypes across cancer types.** DiffDriver identified 50 gene–immune subtype pairs exhibiting significant differential selection across TCGA cancer types. **α**, estimated effect size from DiffDriver (positive values indicate a positive association with the target immune subtype); ***p* value**, nominal *P* value; **FDR**, false discovery rate-adjusted *P* value (*q*-value); **# E = 1**, observed mutation count in the target immune subtype group; **# E = 0**, observed mutation count in the reference group; **E1**, total sample size of the target immune subtype; **E0**, total sample size of the reference group; **Tumor**, TCGA cancer type; **Context**, specific immune subtype (C1–C4) tested as the target phenotype. Only pairs with FDR < 0.1 are shown.

| Gene | α | *p* value | FDR | # E = 1 | # E = 0 | E1 | E0 | Tumor | Context |
|---|---|---|---|---|---|---|---|---|---|
| *BAP1* | 2.45 | 2.71E-02 | 9.47E-02 | 7 | 3 | 129 | 218 | LIHC | C3 |
| *CASP8* | -2.68 | 2.09E-03 | 2.30E-02 | 4 | 41 | 122 | 361 | HNSC | C1 |
| *CASP8* | 2.29 | 5.98E-03 | 5.62E-02 | 40 | 5 | 355 | 128 | HNSC | C2 |
| *CBFB* | 2.74 | 3.17E-03 | 1.98E-02 | 6 | 6 | 163 | 767 | BRCA | C3 |
| *CDH1* | -2.46 | 1.75E-03 | 1.97E-02 | 10 | 40 | 336 | 594 | BRCA | C2 |
| *CDH1* | 2.86 | 1.01E-03 | 8.48E-03 | 17 | 33 | 163 | 767 | BRCA | C3 |
| *CTCF* | -3.36 | 3.15E-03 | 1.97E-02 | 1 | 13 | 336 | 594 | BRCA | C2 |
| *CTNNB1* | -2.62 | 5.81E-03 | 3.48E-02 | 20 | 71 | 129 | 218 | LIHC | C3 |
| *CTNNB1* | 3.02 | 4.19E-04 | 5.87E-03 | 58 | 33 | 153 | 194 | LIHC | C4 |
| *CTNNB1* | 1.02 | 3.74E-03 | 7.49E-02 | 46 | 22 | 107 | 116 | UCEC | C1 |
| *CTNNB1* | -1.72 | 3.65E-05 | 1.19E-03 | 8 | 60 | 71 | 152 | UCEC | C2 |
| *CUL3* | -2.77 | 1.88E-02 | 8.85E-02 | 6 | 9 | 355 | 128 | HNSC | C2 |
| *CUL3* | -1.93 | 4.53E-02 | 9.05E-02 | 4 | 5 | 197 | 73 | KIRP | C3 |
| *EP300* | -4.04 | 6.01E-04 | 9.92E-03 | 2 | 31 | 122 | 361 | HNSC | C1 |
| *EP300* | 4.16 | 4.11E-04 | 6.78E-03 | 31 | 2 | 355 | 128 | HNSC | C2 |
| *FBXW7* | 2.62 | 3.05E-02 | 9.52E-02 | 7 | 2 | 336 | 594 | BRCA | C2 |
| *FGFR3* | -1.59 | 7.27E-05 | 4.58E-03 | 11 | 47 | 160 | 216 | BLCA | C2 |
| *FGFR3* | 1.96 | 2.92E-04 | 1.84E-02 | 15 | 43 | 32 | 344 | BLCA | C4 |

| | | | | | | | | | |
|---|---|---|---|---|---|---|---|---|---|
| *FOXA1* | -3.13 | 2.70E-03 | 1.97E-02 | 1 | 14 | 336 | 594 | BRCA | C2 |
| *FOXA1* | -3.28 | 3.30E-03 | 4.62E-02 | 2 | 6 | 302 | 95 | PRAD | C3 |
| *GATA3* | -3.35 | 1.19E-02 | 5.96E-02 | 0 | 8 | 336 | 594 | BRCA | C2 |
| *GPATCH8* | -3.51 | 1.79E-02 | 8.85E-02 | 2 | 6 | 355 | 128 | HNSC | C2 |
| *HLA-B* | -2.80 | 7.77E-03 | 6.41E-02 | 0 | 15 | 122 | 361 | HNSC | C1 |
| *HLA-B* | 2.87 | 6.81E-03 | 5.62E-02 | 15 | 0 | 355 | 128 | HNSC | C2 |
| *HRAS* | -2.40 | 3.06E-04 | 9.92E-03 | 1 | 31 | 122 | 361 | HNSC | C1 |
| *HRAS* | 2.43 | 2.31E-04 | 6.78E-03 | 31 | 1 | 355 | 128 | HNSC | C2 |
| *KIT* | -4.65 | 2.19E-05 | 6.56E-05 | 0 | 24 | 41 | 106 | TGCT | C1 |
| *KIT* | 5.04 | 1.74E-05 | 5.22E-05 | 24 | 0 | 103 | 44 | TGCT | C2 |
| *KRAS* | 2.38 | 1.34E-03 | 4.21E-02 | 6 | 7 | 32 | 344 | BLCA | C4 |
| *KRAS* | -3.10 | 2.88E-02 | 9.52E-02 | 0 | 5 | 336 | 594 | BRCA | C2 |
| *KRAS* | 4.32 | 7.12E-03 | 2.85E-02 | 1 | 4 | 3 | 267 | KIRP | C1 |
| *KRAS* | -1.38 | 4.32E-03 | 8.20E-02 | 29 | 102 | 132 | 287 | LUAD | C2 |
| *KRAS* | 1.96 | 2.78E-04 | 5.29E-03 | 63 | 68 | 169 | 250 | LUAD | C3 |
| *KRAS* | -3.03 | 4.23E-03 | 6.34E-03 | 1 | 16 | 41 | 106 | TGCT | C1 |
| *KRAS* | 3.17 | 4.67E-03 | 7.01E-03 | 16 | 1 | 103 | 44 | TGCT | C2 |
| *MAP3K1* | -1.19 | 2.83E-02 | 9.52E-02 | 5 | 21 | 336 | 594 | BRCA | C2 |
| *MET* | 2.02 | 3.25E-02 | 9.05E-02 | 19 | 1 | 197 | 73 | KIRP | C3 |
| *NFE2L2* | 2.12 | 7.46E-03 | 3.48E-02 | 8 | 3 | 129 | 218 | LIHC | C3 |
| *NRAS* | -2.83 | 4.32E-02 | 4.32E-02 | 0 | 6 | 41 | 106 | TGCT | C1 |
| *NRAS* | 2.88 | 3.91E-02 | 3.91E-02 | 6 | 0 | 103 | 44 | TGCT | C2 |
| *PIK3CA* | -1.75 | 1.40E-04 | 3.49E-03 | 96 | 222 | 336 | 594 | BRCA | C2 |
| *PIK3CA* | 2.13 | 1.02E-03 | 8.48E-03 | 73 | 245 | 163 | 767 | BRCA | C3 |

| | | | | | | | | | |
|---|---|---|---|---|---|---|---|---|---|
| *SMARCA4* | 2.34 | 1.20E-02 | 7.90E-02 | 10 | 9 | 122 | 361 | HNSC | C1 |
| *SMARCA4* | -2.23 | 1.49E-02 | 8.85E-02 | 9 | 10 | 355 | 128 | HNSC | C2 |
| *SMARCA4* | -2.78 | 4.77E-03 | 4.54E-02 | 3 | 29 | 169 | 250 | LUAD | C3 |
| *TP53* | -5.23 | 6.45E-07 | 1.61E-05 | 8 | 219 | 163 | 767 | BRCA | C3 |
| *TP53* | -3.90 | 1.07E-04 | 1.50E-03 | 14 | 72 | 129 | 218 | LIHC | C3 |
| *TP53* | -1.22 | 1.96E-03 | 7.49E-02 | 16 | 37 | 107 | 116 | UCEC | C1 |
| *TP53* | 1.58 | 5.95E-05 | 1.19E-03 | 31 | 22 | 71 | 152 | UCEC | C2 |

**Supplementary Table 5 |** Pan-cancer differential selection of driver genes across immune subtypes. The 20 gene–immune-subtype associations significant at FDR < 0.1 from the pan-cancer analysis (8,253 samples across 27 TCGA cancer types; 220 recurrently mutated driver genes; subtypes C1–C4, one-vs-rest, adjusting for age, sex, and tumor type). α is the subtype coefficient (positive: stronger selection within the subtype; negative: stronger selection in the remaining subtypes).

| Gene | Immune subtype | Direction | α | p value | FDR | Mut freq (%) | # tumor types | # samples |
|---|---|---|---|---|---|---|---|---|
| *TP53* | C2 | up in subtype | 1.74 | 1.13e-16 | 0.000 | 40.6 | 23 | 7153 |
| *KRAS* | C2 | up in rest | -1.00 | 2.11e-05 | 0.002 | 12.8 | 14 | 4835 |
| *JAK2* | C2 | up in subtype | 3.10 | 2.43e-04 | 0.018 | 1.4 | 6 | 3006 |
| *COL5A1* | C2 | up in subtype | 0.83 | 6.20e-04 | 0.032 | 3.1 | 16 | 5843 |
| *CASP8* | C2 | up in subtype | 0.92 | 7.28e-04 | 0.032 | 3.3 | 8 | 3784 |
| *HLA-A* | C2 | up in subtype | 1.99 | 1.97e-03 | 0.072 | 2.4 | 4 | 1547 |
| *PPP2R1A* | C2 | up in subtype | 1.49 | 2.64e-03 | 0.079 | 6.0 | 5 | 1595 |
| *HLA-B* | C2 | up in subtype | 1.14 | 3.23e-03 | 0.079 | 2.0 | 7 | 2612 |
| *EZH2* | C2 | up in subtype | 2.97 | 3.57e-03 | 0.079 | 1.3 | 4 | 1686 |
| *RB1* | C2 | up in subtype | 0.85 | 3.59e-03 | 0.079 | 4.4 | 11 | 4259 |
| *FAT1* | C2 | up in subtype | 0.54 | 4.85e-03 | 0.097 | 6.1 | 15 | 6084 |
| *TP53* | C3 | up in rest | -2.41 | 4.17e-29 | 0.000 | 40.6 | 23 | 7153 |

| | | | | | | | | |
|---|---|---|---|---|---|---|---|---|
| *WHSC1* | C3 | up in rest | -2.78 | 3.96e-04 | 0.044 | 1.6 | 7 | 3412 |
| *FAT1* | C3 | up in rest | -1.06 | 6.09e-04 | 0.045 | 6.1 | 15 | 6084 |
| *SMARCA4* | C3 | up in rest | -1.45 | 1.03e-03 | 0.048 | 3.7 | 15 | 5987 |
| *BRCA1* | C3 | up in rest | -2.11 | 1.20e-03 | 0.048 | 2.6 | 10 | 4308 |
| *LATS2* | C3 | up in rest | -3.36 | 1.30e-03 | 0.048 | 1.8 | 6 | 2271 |
| *POLQ* | C3 | up in rest | -1.19 | 2.81e-03 | 0.088 | 3.6 | 14 | 5547 |
| *CIC* | C4 | up in rest | -2.60 | 6.47e-06 | 0.001 | 3.9 | 10 | 4096 |
| *EGFR* | C4 | up in subtype | 1.66 | 7.67e-05 | 0.008 | 3.8 | 12 | 5023 |

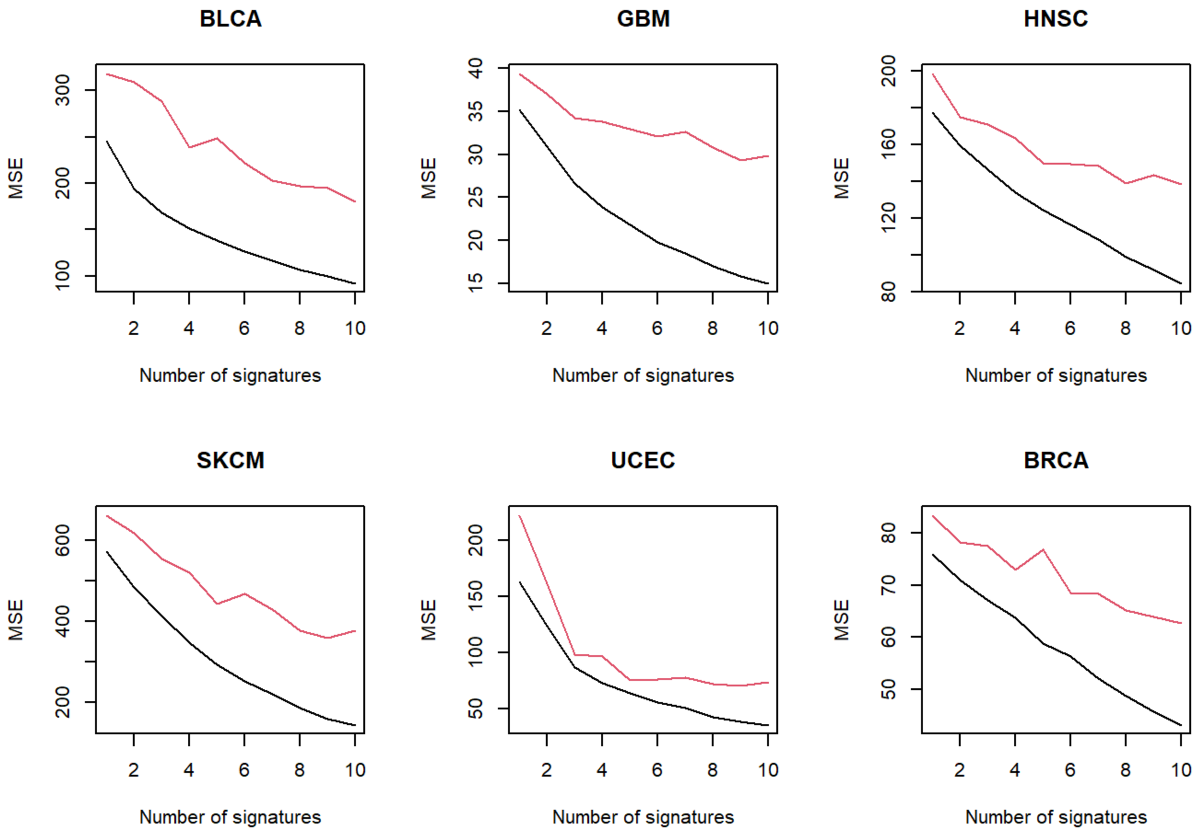


**Supplementary Figure 1. Background Mutation Model (BMM) fitting with varying number of signatures.** We fit the synonymous mutation data using the DiffDriver BMM or NMF via the "Lee and Seung method"[2]. The NMF method is commonly used in existing literature to extract mutational signatures[3]. The DiffDriver BMM is essentially fitting a Poisson NMF model. The data to train the model here is a matrix of mutation count for each mutation type (96 in total) in each individual. We calculated mean square error (MSE) for the DiffDriver BMM (black line) and the NMF model (red line). Results for 6 representative tumor types are shown.

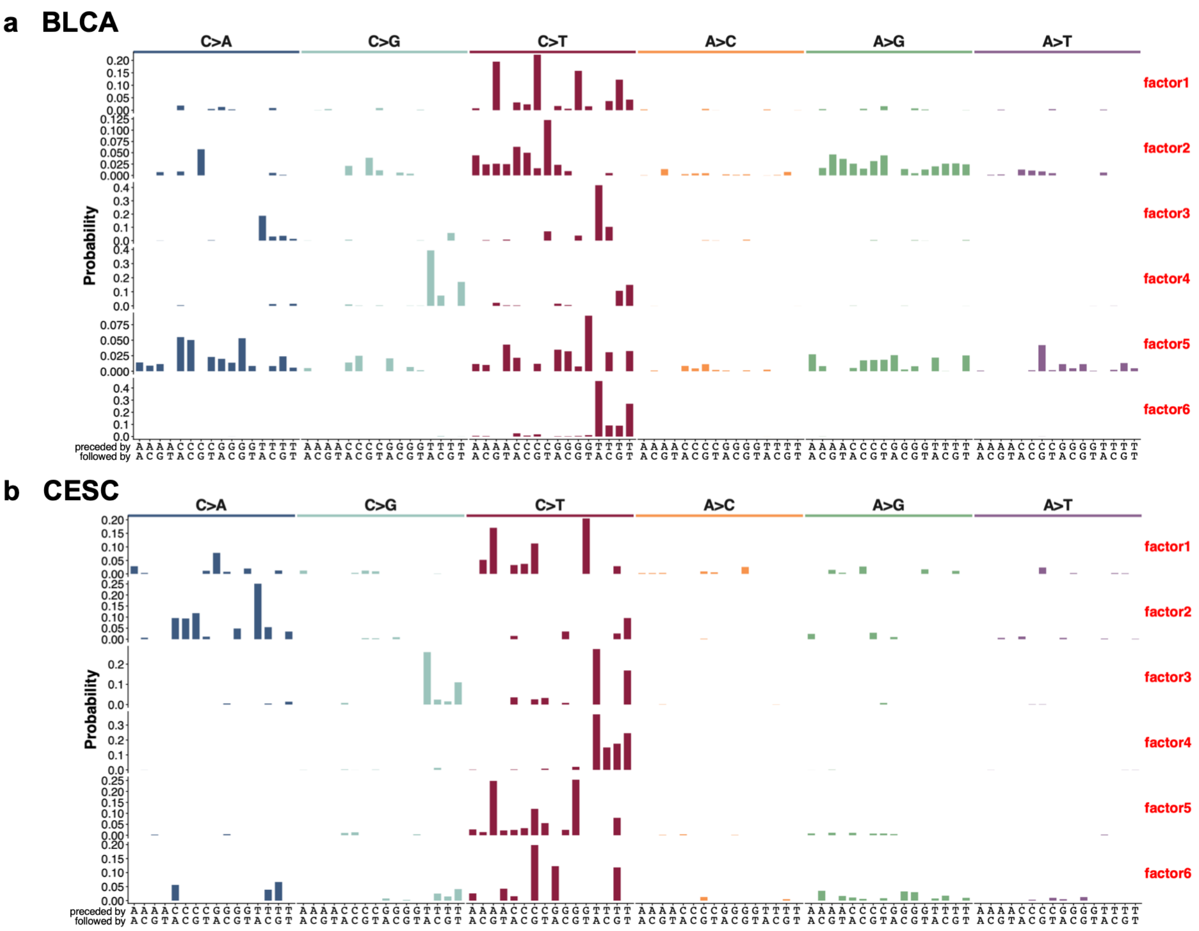


**Supplementary Figure 2. DiffDriver captures APOBEC3-associated mutational signatures in TCGA bladder and cervical cancer cohorts.** Mutational signatures (Factor 1–6) extracted by the DiffDriver Background Mutation Model (BMM) from The Cancer Genome Atlas (TCGA) bladder urothelial carcinoma and cervical squamous cell carcinoma cohorts. **a, BLCA cohort; b, CESC cohort.** Each profile displays the probability distribution of base substitutions across 96 trinucleotide contexts. The contexts are categorized into six major mutation subtypes (e.g., C>A, C>T), indicated by the colored bars at the top. The x-axis indicates the specific sequence motif, defined by the mutated base and its 5′ (top row) and 3′ (bottom row) flanking bases. The y-axis represents the contribution of each mutation type to the signature.

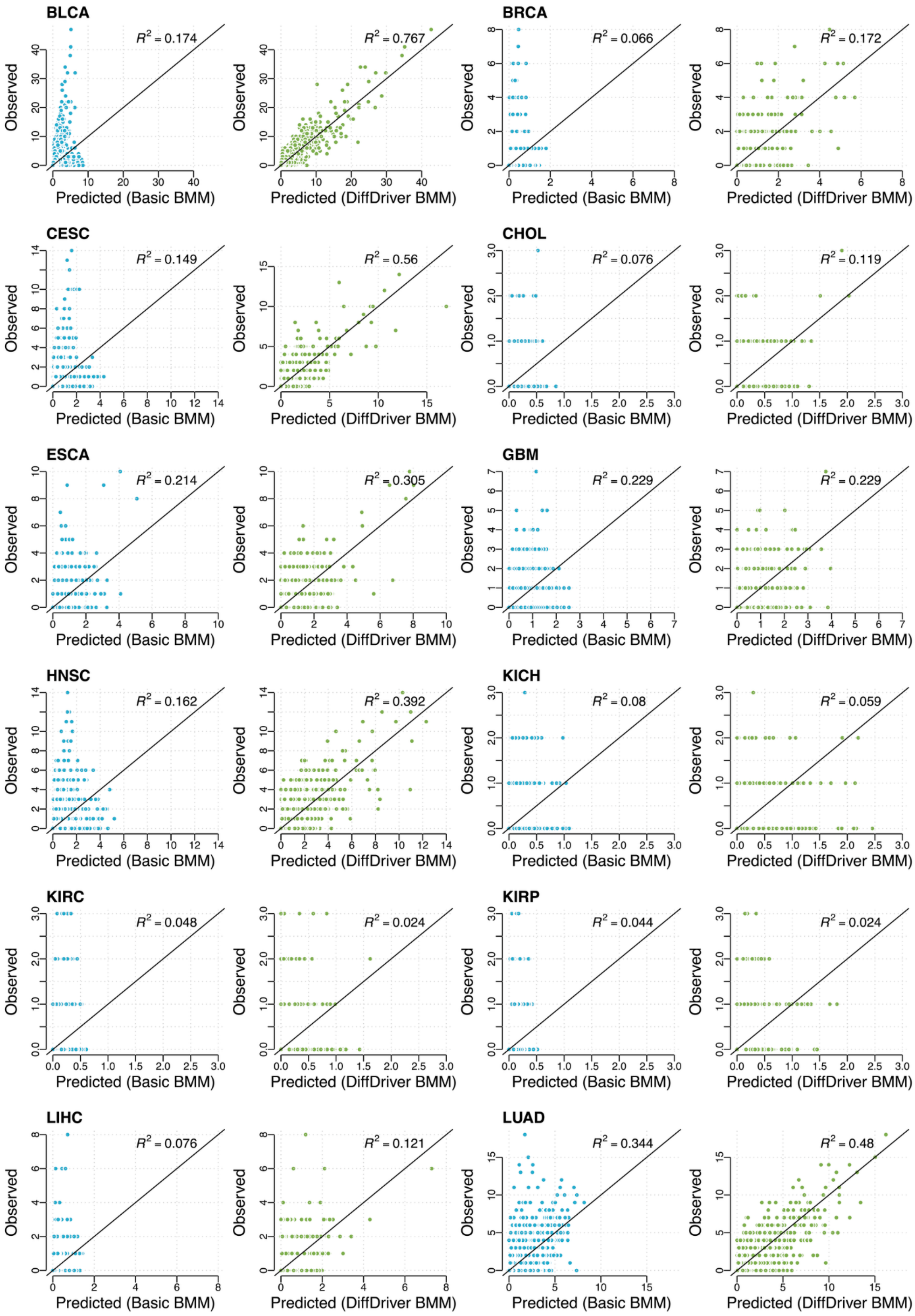
BLCA
R2 = 0.174
R2 = 0.767
BRCA
R2 = 0.066
R2 = 0.172
CESC
R2 = 0.149
R2 = 0.56
CHOL
R2 = 0.076
R2 = 0.119
ESCA
R2 = 0.214
R2 = 0.305
GBM
R2 = 0.229
R2 = 0.229
HNSC
R2 = 0.162
R2 = 0.392
KICH
R2 = 0.08
R2 = 0.059
KIRC
R2 = 0.048
R2 = 0.024
KIRP
R2 = 0.044
R2 = 0.024
LIHC
R2 = 0.076
R2 = 0.121
LUAD
R2 = 0.344
R2 = 0.48
Observed
Predicted (Basic BMM)
Predicted (DiffDriver BMM)

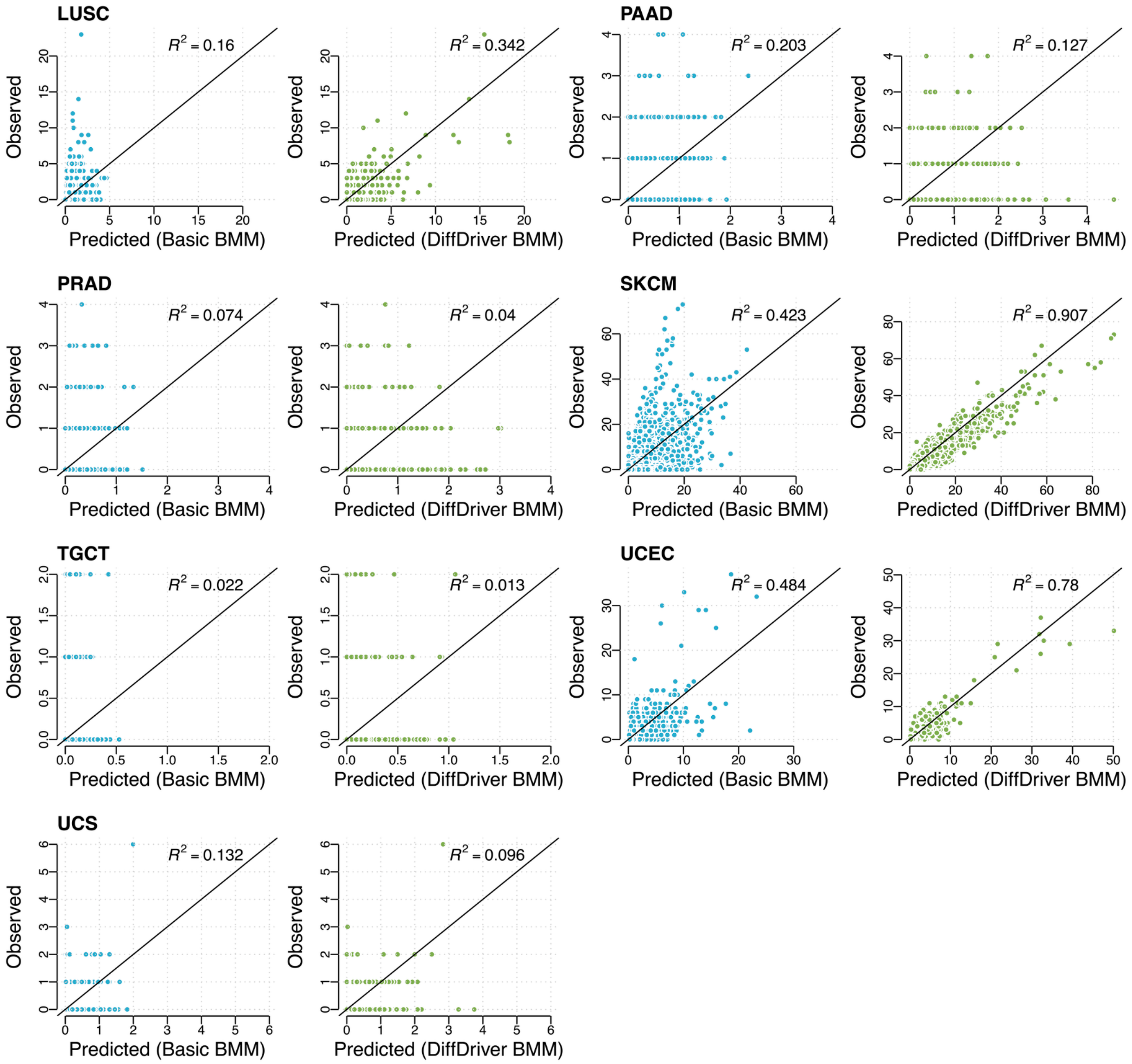


**Supplementary Figure 3. Calibration of background mutation models across 19 cancer types.** This figure displays goodness-of-fit scatter plots across 19 cancer types, plotting the observed number of mutations (y-axis) against the predicted number (x-axis) for 96 trinucleotide contexts. The comparison highlights predictions generated by the Basic BMM (left), formulated under the assumption of a uniform mutation rate across individuals, versus those generated by the DiffDriver BMM (right), which accounts for patient-specific mutation rates. The solid black line represents the diagonal ($y = x$) indicating perfect calibration, and the specific cancer type along with the coefficient of determination ($R^2$) is annotated within each plot.

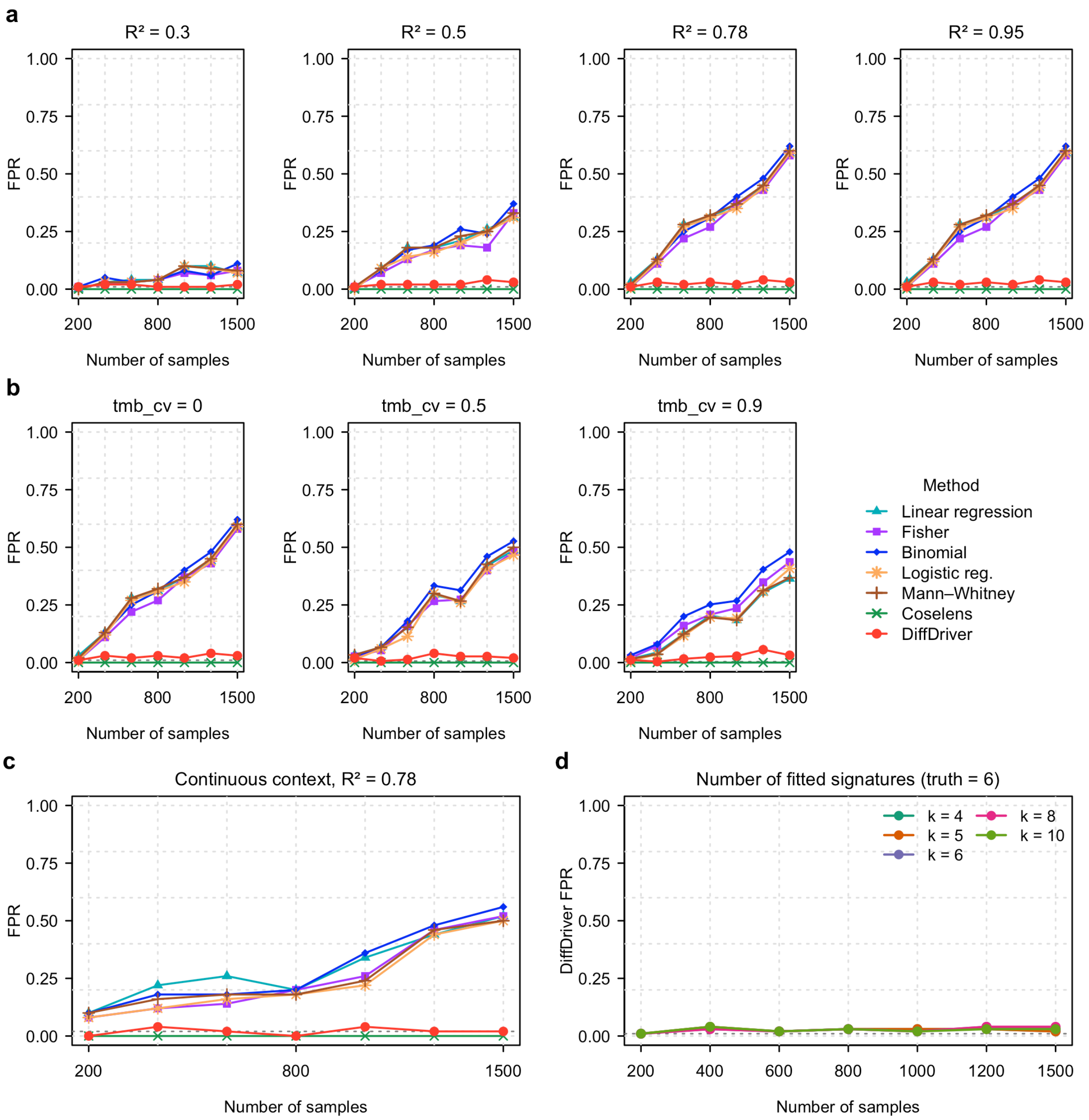


**Supplementary Figure 4. DiffDriver controls false positives under context–signature confounding across a range of simulation assumptions.** a–d, In a controlled confounded-null simulation—mutations are simulated for a single gene (*ERBB3*) whose hotspots fall in the peak trinucleotide context of LUAD Signature 3, and the context variable is correlated with that signature but has zero correlation with selection, so every detection is a false positive. We vary the simulation assumptions and plot the false-positive rate (FPR) versus sample size for all seven methods; DiffDriver and Coselens estimate the background from a simulated genome-wide silent (synonymous) mutation matrix, as on real data. In every panel, FPR is assessed at a significance threshold of 1/Niter (dashed line)—the finest level resolvable with Niter iterations—

i.e. 0.01, 0.0067, 0.004, and 0.02 for the 100-, 150-, 250-, and 50-iteration settings, respectively. (a) Confounding strength: as the context–signature correlation $R^2$ increases (0.3→0.95), the FPR of the count-based methods (Linear regression, Fisher's exact test, Binomial test, Logistic regression, Mann–Whitney) rises monotonically (10% to 60%), whereas DiffDriver and Coselens remain controlled. (b) Mutation-burden heterogeneity: inflating each sample's total burden by a log-normal factor (tmb_cv = 0, 0.5, 0.9; up to a ≈5–10× spread) leaves the picture unchanged, because the confounding resides in signature composition rather than the per-sample total. (c) A continuous confounded context ($R^2$ = 0.78) still misleads the count-based methods but not DiffDriver or Coselens. (d) Number of fitted signatures: refitting the silent matrix with k ∈ {4, 5, 6, 8, 10} (truth = 6) leaves DiffDriver's false-positive control essentially unchanged (≈1–4%). Sample sizes N = 200–1,500; 100 iterations per point (150 and 250 for the two variable-burden panels in b, 50 for the continuous panel in c).

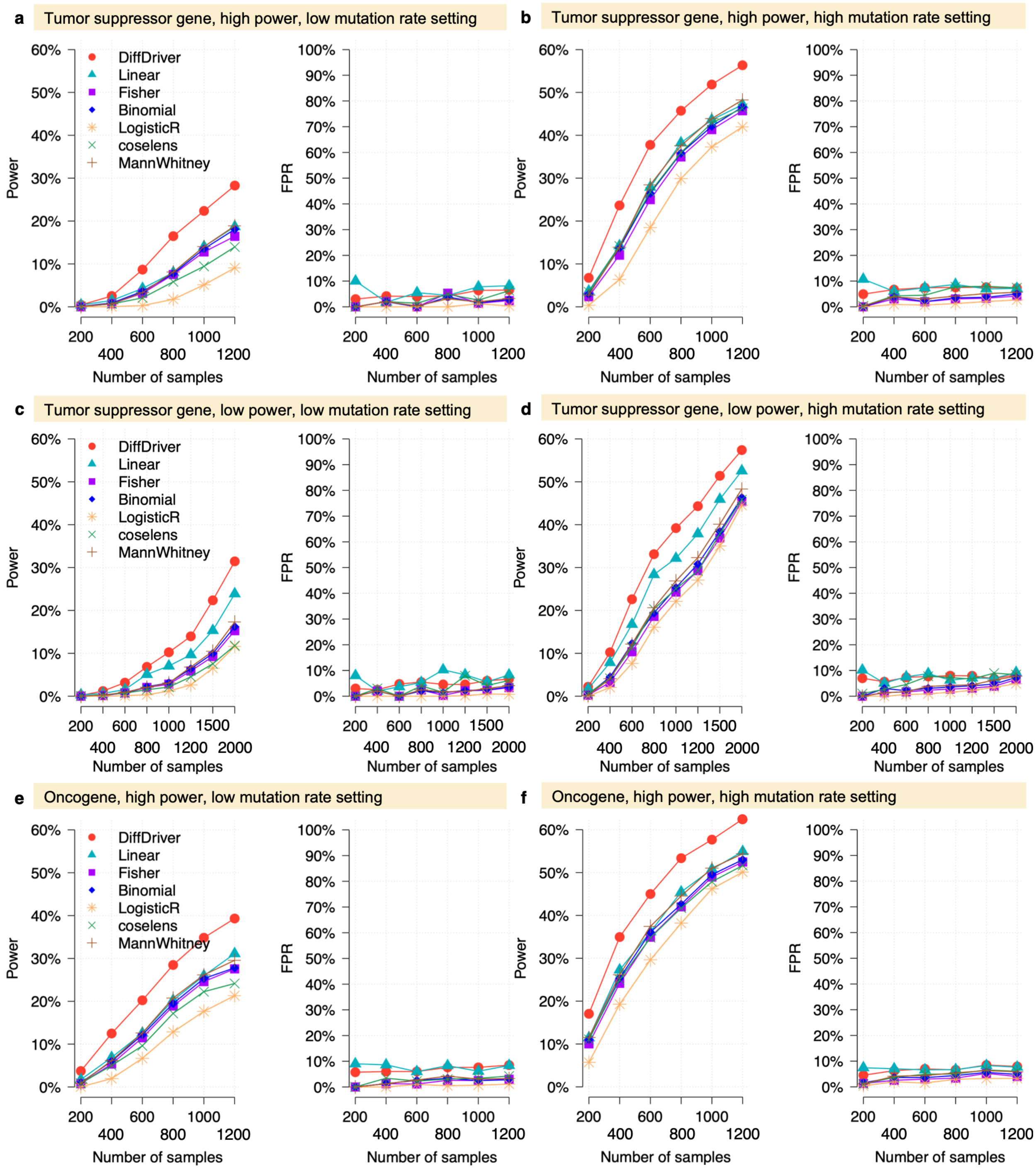


**Supplementary Figure 5. Benchmarking DiffDriver performance against standard methods using simulated data with a continuous context. a-f,** Performance comparison of DiffDriver (red) with six baseline methods (Linear regression, Fisher's exact test, Binomial test, Logistic regression, Coselens, and the Mann-Whitney test) in detecting driver genes across increasing sample sizes. The simulations evaluate performance for (a–d) tumor suppressor genes under high-power (a, b) and low-power (c, d) settings, and for (e, f) oncogenes under high-power settings. Panels are further stratified by low (a, c, e) and high (b, d, f) background mutation rates. In each panel, the left plot shows the statistical power, and the right plot indicates the False Positive Rate (FPR).

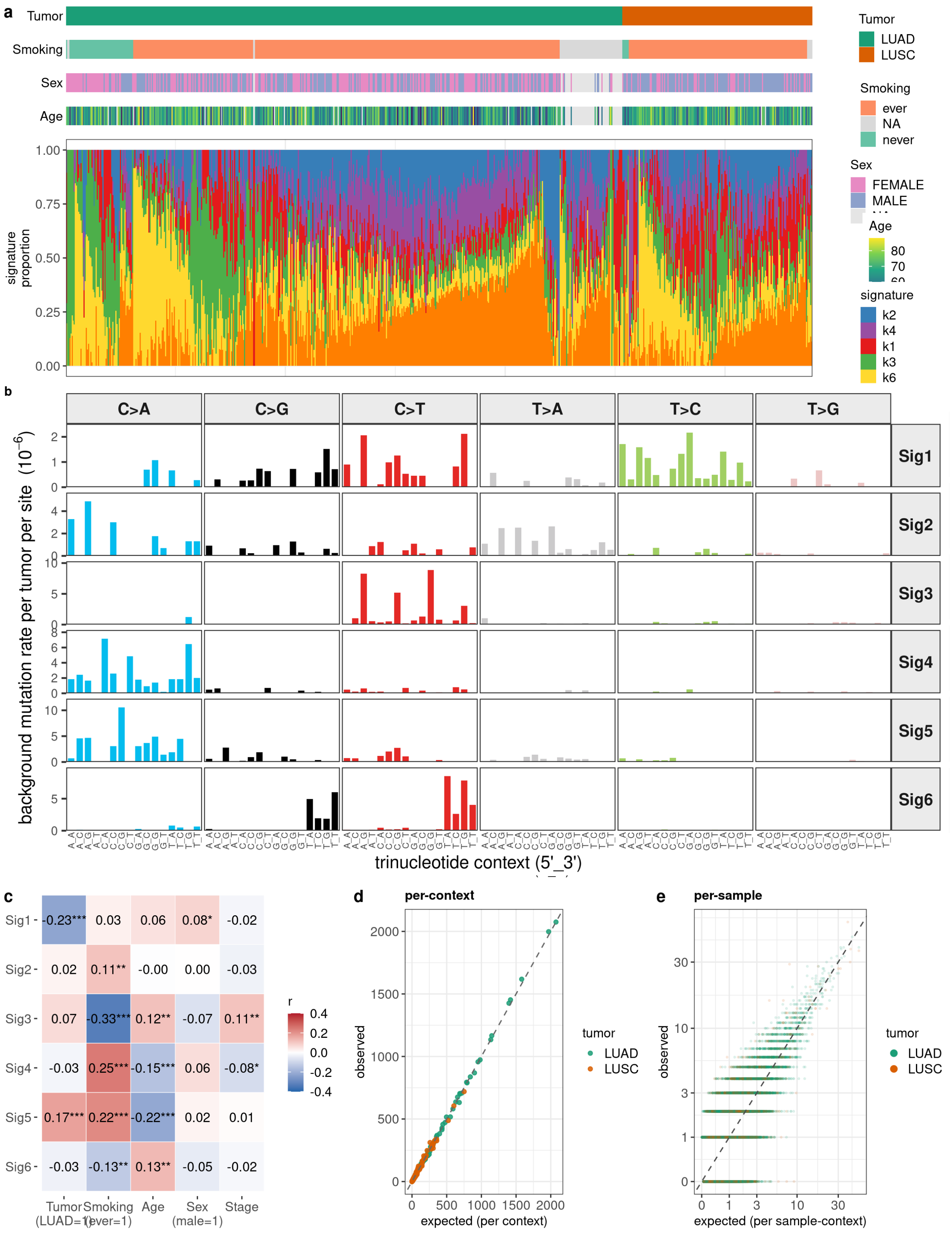


**Supplementary Figure 6. The DiffDriver signature-based background model captures sample-level characteristics and mutation-rate heterogeneity in lung cancer.** Analysis of

704 TCGA lung tumors (530 lung adenocarcinoma, LUAD; 174 lung squamous cell carcinoma, LUSC) using the six-signature ($k = 6$) background mutation-rate model estimated by DiffDriver. **a**, Structure plot of the per-sample mutational-signature loadings (679 samples with ≥5 mutations). Samples are grouped by tumor type and then smoking status, and ordered within each group by signature similarity; the tracks above show each sample's tumor type, smoking status (ever/never), sex, and age. **b**, The six inferred signatures across the 96 trinucleotide contexts (standard single-base-substitution layout; columns, the six substitution classes; rows, signatures; x-axis, 5′ and 3′ flanking bases), shown on the actual per-site background-mutation-rate scale: bars give the modeled background mutation rate per tumor per site contributed by each signature (units of $10^{-6}$; y-axis scaled per signature). Even the most active contexts contribute only a few × $10^{-6}$ per tumor, so a site recurrently mutated across many tumors reflects selection rather than background. Signature 4 is dominated by C>A transversions (COSMIC SBS4, tobacco smoking) and Signature 3 by C>T transitions at CpG (SBS1, clock-like aging). **c**, Pearson correlation between each sample-level covariate (tumor type, smoking status, age, sex, stage) and each signature loading; asterisks denote significance (*$P<0.05$, **$P<0.01$, ***$P<0.001$). **d, e**, Goodness of fit of the signature model, comparing model-expected to observed mutation counts, summed per trinucleotide context (d) and at the individual sample-by-context level (e); points are colored by tumor type and the dashed line is $y = x$.

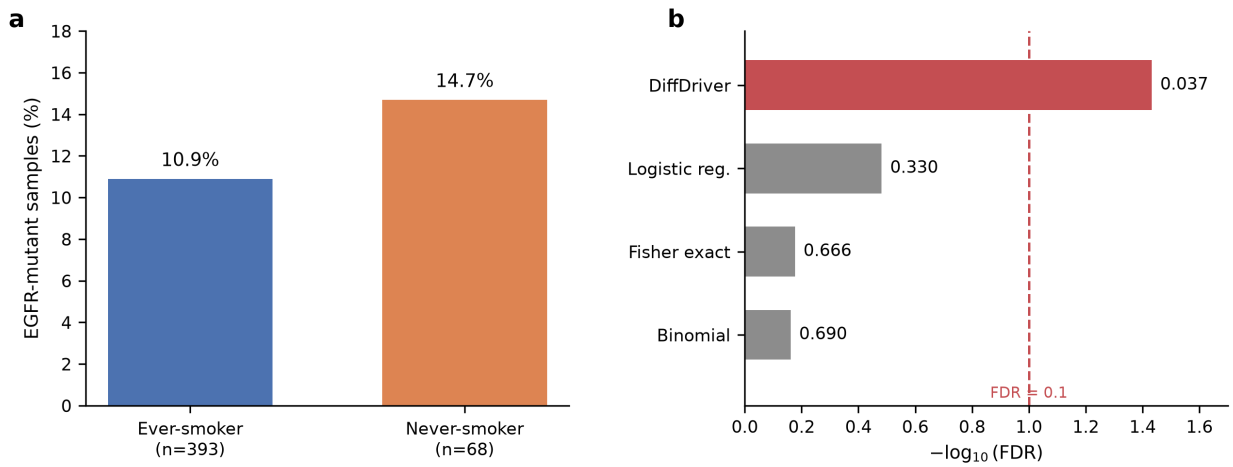


**Supplementary Figure 7. DiffDriver detects context-associated selection of EGFR in never-smoker lung adenocarcinoma that frequency-based methods miss.** EGFR mutation status in TCGA LUAD stratified by smoking history (393 ever-smokers, 68 never-smokers). **a**, the frequency of EGFR-mutant samples is similar between ever- and never-smokers (10.9% vs 14.7%; Fisher's exact $p$ = 0.42, not significant). **b**, when EGFR is tested for smoking-differential selection, only DiffDriver identifies significantly stronger selection in never-smokers (FDR = 0.037), whereas frequency-based methods (logistic regression, Fisher's exact test, binomial test) do not (FDR > 0.3). The dashed vertical line marks FDR = 0.1.

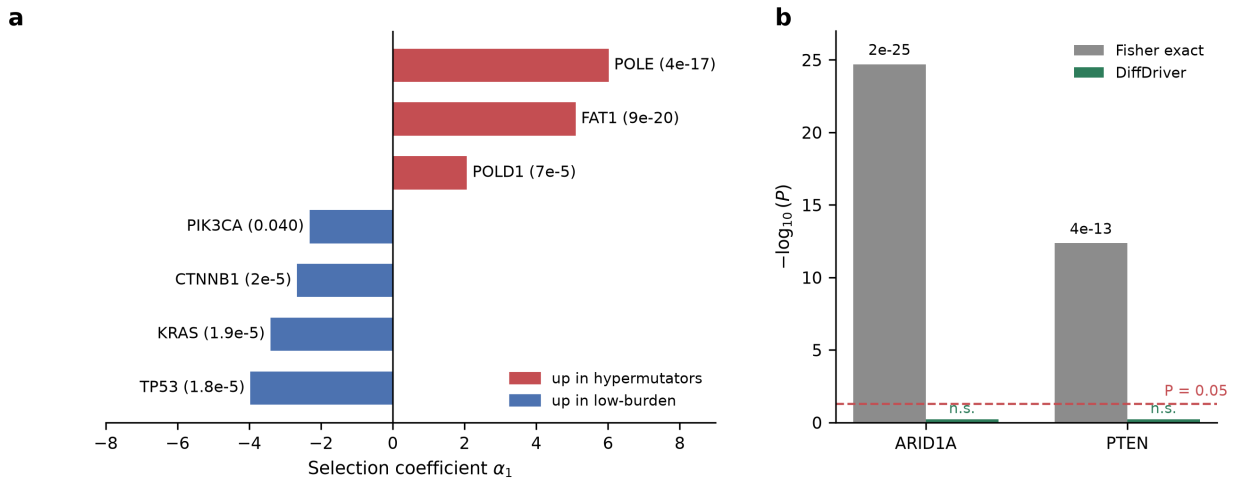


**Supplementary Figure 8. DiffDriver distinguishes differential selection from mutation-rate heterogeneity in hypermutated endometrial carcinoma.** DiffDriver was applied to TCGA UCEC (512 samples: 74 hypermutated tumors with >1,000 mutations and 438 low-burden tumors), testing UCEC driver genes together with POLE and POLD1 for differential selection by mutation burden, adjusting for age. **a**, Selection coefficient ($\alpha_1$) for each significant gene (FDR < 0.05); positive values indicate stronger selection in hypermutated tumors (red) and negative values stronger selection in low-burden tumors (blue). POLE (FDR = $4\times10^{-17}$) and POLD1, whose proofreading-domain mutations cause the ultramutator phenotype, are the strongest hypermutator-associated genes and serve as positive controls, along with FAT1; TP53, KRAS, CTNNB1, and PIK3CA—canonical drivers of the non-hypermutated subtypes—are under stronger selection in low-burden tumors. **b**, ARID1A and PTEN show large differences in raw mutation frequency between the two groups (Fisher's exact P = $2\times10^{-25}$ and $4\times10^{-13}$, higher frequency in hypermutators) but are correctly not called as differentially selected by DiffDriver (n.s.), because their elevated frequencies in hypermutated tumors are explained by the higher background mutation rate rather than by stronger selection.

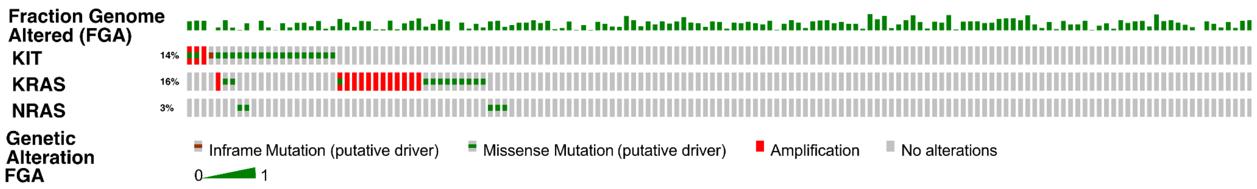


**Supplementary Figure 9. Landscape of genomic alterations in *KIT*, *KRAS*, and *NRAS*.** The Oncoprint visualization depicts the distribution of somatic alterations across the study cohort. Each column represents an individual sample. The top histogram tracks the Fraction of Genome Altered (FGA), serving as a proxy for chromosomal instability. Genetic alterations are color-coded by type: amplifications (red), putative driver missense mutations (green), and inframe mutations (brown). Grey bars indicate samples with no detected alterations in the specified genes. The percentages on the left indicate the overall alteration frequency of each gene within the cohort.

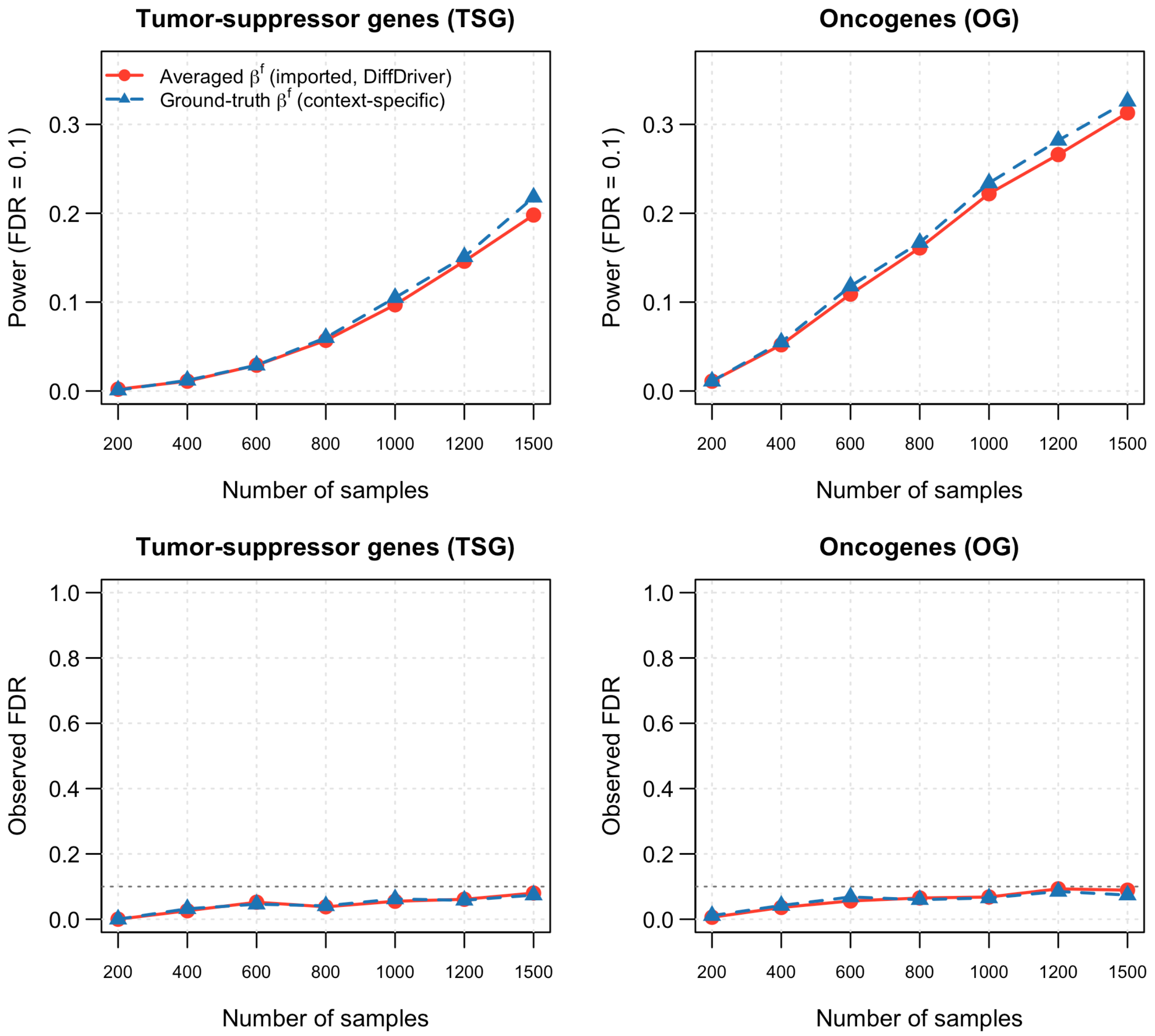


**Supplementary Figure 10. DiffDriver's power is insensitive to the use of imported, averaged functional effect sizes ($\beta^f$).** In the power simulation, mutations are generated under the true, context-specific $\beta^f$, and DiffDriver is run on the identical simulated cohorts using either the imported/averaged $\beta^f$ (as in all main-text results; pre-estimated by DriverMAPS) or the true generating $\beta^f$ (the upper bound achievable by perfectly re-estimating $\beta^f$ for each context). Top: statistical power (fraction of the 50 signal genes detected at a target FDR of 0.1); bottom: observed false discovery rate (from the 150 null genes), each versus sample size, for tumor-suppressor genes (left) and oncogenes (right). Although the imported $\beta^f$ are attenuated relative to the truth, power differs by ≤2 percentage points at every sample size and the FDR remains controlled at the nominal 10%, indicating that the imported effect sizes do not materially dampen DiffDriver's power to detect context-specific differential selection.
